# A new domestic cat genome assembly based on long sequence reads empowers feline genomic medicine and identifies a novel gene for dwarfism

**DOI:** 10.1101/2020.01.06.896258

**Authors:** Reuben M. Buckley, Brian W. Davis, Wesley A. Brashear, Fabiana H. G. Farias, Kei Kuroki, Tina Graves, LaDeana W. Hillier, Milinn Kremitzki, Gang Li, Rondo Middleton, Patrick Minx, Chad Tomlinson, Leslie A. Lyons, William J. Murphy, Wesley C. Warren

**Affiliations:** Department of Veterinary Medicine and Surgery, College of Veterinary Medicine, University of Missouri, Columbia, Missouri 65211, USA; Department of Veterinary Integrative Biosciences, Interdisciplinary Program in Genetics, College of Veterinary Medicine, Texas A&M University, College Station, Texas 77843, USA; McDonnell Genome Institute, Washington University, School of Medicine, St Louis, Missouri 63108, USA; Veterinary Medical Diagnostic Laboratory, College of Veterinary Medicine, University of Missouri, Columbia, Missouri, 65211, USA; Nestlé Purina Research, Saint Louis, Missouri 63164, USA; Division of Animal Sciences, School of Medicine, University of Missouri, Columbia, Missouri 65211, USA

**Keywords:** animal model, chondrodysplasia, dwarfism, *Felis catus*, long-read, Precision Medicine

## Abstract

The domestic cat (*Felis catus*) numbers over 94 million in the USA alone, occupies households as a companion animal, and, like humans, suffers from cancer and common and rare diseases. However, genome-wide sequence variant information is limited for this species. To empower trait analyses, a new cat genome reference assembly was developed from PacBio long sequence reads that significantly improve sequence representation and assembly contiguity. The whole genome sequences of 54 domestic cats were aligned to the reference to identify single nucleotide variants (SNVs) and structural variants (SVs). Across all cats, 16 SNVs predicted to have deleterious impacts and in a singleton state were identified as high priority candidates for causative mutations. One candidate was a stop gain in the tumor suppressor *FBXW7*. The SNV is found in cats segregating for feline mediastinal lymphoma and is a candidate for inherited cancer susceptibility. SV analysis revealed a complex deletion coupled with a nearby potential duplication event that was shared privately across three unrelated dwarfism cats and is found within a known dwarfism associated region on cat chromosome B1. This SV interrupted *UDP-glucose 6-dehydrogenase (UGDH)*, a gene involved in the biosynthesis of glycosaminoglycans. Importantly, *UGDH* has not yet been associated with human dwarfism and should be screened in undiagnosed patients. The new high-quality cat genome reference and the compilation of sequence variation demonstrate the importance of these resources when searching for disease causative alleles in the domestic cat and for identification of feline biomedical models.

**Author summary:** The practice of genomic medicine is predicated on the availability of a high quality reference genome and an understanding of the impact of genome variation. Such resources have lead to countless discoveries in humans, however by working exclusively within the framework of human genetics, our potential for understanding diseases biology is limited, as similar analyses in other species have often lead to novel insights. The generation of Felis_catus_9.0, a new high quality reference genome for the domestic cat, helps facilitate the expansion of genomic medicine into the *felis* lineage. Using Felis_catus_9.0 we analyze the landscape of genomic variation from a collection of 54 cats within the context of human gene constraint. The distribution of variant impacts in cats is correlated with patterns of gene constraint in humans, indicating the utility of this reference for identifying novel mutations that cause phenotypes relevant to human and cat health. Moreover, structural variant analysis revealed a novel variant for feline dwarfism in *UGDH*, a gene that has not been associated with dwarfism in any other species, suggesting a role for *UGDH* in cases of undiagnosed dwarfism in humans.

## Background

In the veterinary clinic, the practice of genomic medicine is impending [1]. With actionable genetic information in hand, companion animal therapeutic interventions are feasible, including treatment of animal patients prior to, or to prevent the appearance of, more severe symptoms and allow therapeutic administration of drugs with higher efficacy and fewer side effects. Genomic information can also alert veterinarians to imminent disease risks for diagnostic consideration. Each of these applications could significantly enhance veterinary medicine, however, none are currently in practice. As in human medicine, formidable challenges exist for the implementation of genomic-based medicine, including accurate annotation of the genome and databases of genetic variation from well-phenotyped individuals that include both single nucleotide variant (SNV) and structural variant (SV) discovery and annotation [2-4]. Targeted individual companion animal genome information is becoming more readily available, cost effective, and tentatively linked to the actionable phenotypes via direct-to-consumer DNA testing. Thus correct interpretation of DNA variants is of the upmost importance for communicating findings to clinicians practicing companion animal genomic medicine [5].

Companion animals suffer from many of the same diseases as humans, with over 600 different phenotypes identified as comparative models for human physiology, biology, development, and disease [6, 7]. In domestic cats, at least 70 genes are shown to harbor single and multiple DNA variants that are associated with disease [8] with many more discoveries expected as health care improves.. The genetic and clinical manifestations of most of these known variants are described in the Online Mendelian Inheritance in Animals [6]. Examples include common human diseases such as cardiomyopathy [9], retinal degenerations [10], and polycystic kidney disease [11]. Veterinarians, geneticists and other researchers are actively banking DNA from companion animals and attempting to implement genomic medicine [1]. Once coupled with the quickly advancing sequencing technology and exploitable results, genomic medicine in companion animals promises to expand the comparative knowledge of mechanisms of action across species. Despite the continuing successful discovery of feline disease variants using both candidate gene and whole genome sequencing (WGS) approaches [1, 12-15], the understanding of normal and disease sequence variation in the domestic cat and interrogation of gene structure and function is limited by an incomplete genome assembly.

A fundamental hurdle hampering the interpretation of feline disease variant data is the availability of a high-quality, gapless reference genome. The previous domestic cat reference, Felis_catus_8.0, contains over 300,000 gaps, compromising its utility for identifying all types of sequence variation [16], in particular SVs. In conjunction with various mapping technologies, such as optical resolved physical maps, recent advances in the use of long-read sequencing and assembly technology has produced a more complete genome representation (i.e., fewer gaps) for many species [17-20].

Another hurdle for performing feline genomic medicine is the availability of WGS data from various breeds of cat with sufficient sequencing depth to uncover rare alleles and complex structural variants. Knowledge of variant frequency and uniqueness among domestic cats is very limited and is crucial in the identification of causal alleles. As a result of the paucity of sequence variant data across breeds, the 99 Lives Cat Genome Sequencing Initiative was founded as a centralized resource with genome sequences produced of similar quality and techniques. The resource supports researchers with variant discovery for evolutionary studies and identifying the genetic origin of inherited diseases and can assist in the development of high-density DNA arrays for complex disease studies in domestic cats [1, 21-24].

Here we present a new version of the domestic cat genome reference (Cinnamon, an Abyssinian breed), generated from deep sequence coverage of long-reads and scaffolding from an optical map (BioNano) and a high-density genetic linkage map [16]. Published cat genomes from the 99 Lives Cat Genome Consortium [1, 23] were aligned to the Felis_catus_9.0 reference to discover a plethora of unknown SNVs and SVs (multi-base insertions and deletions), including a newly identified structural variant (SV) for feline disproportionate dwarfism. Our case study of dwarfism demonstrates when disease phenotypes are coupled with revised gene annotation and sequence variation ascertained from diverse breeds, the new cat genome assembly is a powerful resource for trait discovery. This enables the future practice of feline genomic medicine and improved ascertainment of biomedical models relevant to human health.

## Results

### Genome assembly

A female Abyssinian cat (Cinnamon) was sequenced to high-depth (72-fold coverage) using real-time (SMRT; PacBio) sequence data and all sequence reads were used to generate a *de novo* assembly. Two PacBio instruments were used to produce average read insert lengths of 12 kb (RSII) and 9 kb (Sequel). The ungapped assembly size was 2.48 Gb and is comparable in size to other assembled carnivores (**Table 1**). There were 4,909 total contigs compared to 367,672 contigs in Felis_catus_8.0 showing a significant reduction in sequence gaps. The assembly contiguity metric of N50 contig and scaffold lengths were 42 and 84 Mb, respectively (**Table 1**). The N50 contig length of other PacBio sequenced carnivore assemblies are less contiguous, ranging from 3.13 Mb to 20.91 Mb (**Table 1**). Across carnivores, RepeatMasker showed consistent measures of total interspersed repeat content, (with 43% in Felis_catus*_*9.0; **S1 Table**) [25]. Due to repetitive and other genome architecture features, 1.8% (46 Mb) of all assembled sequences remained unassigned to any chromosome position. These sequences had an N50 scaffold length of 12,618 bp, demonstrating the assembly challenge of some repeat types in diploid genome assemblies, even of an inbred individual.

**Table 1.** Representative assembly metrics for various chromosome level assembled carnivore genomes^1^.

| Assembly | Species | Breed | Isolate | Release date (MM/DD/YY) | Sequencing technology | Genome coverage | Total ungapped length (Gb) | Scaffold N50 (Mb) | Contig N50 (Mb) | Unplaced length (Mb) |
| --- | --- | --- | --- | --- | --- | --- | --- | --- | --- | --- |
| Felis_catus_9.0 | *Felis catus* (domestic cat) | Abyssinian | Cinnamon | 11/20/17 | PacBio; 454 Titanium; Illumina; Sanger dideoxy sequencing | 72x | 2.48 | 83.97 | 41.92 | 46.02 |
| Felis_catus_8.0 | *Felis catus* (domestic cat) | Abyssinian | Cinnamon | 11/07/14 | Sanger; 454 Titanium; Illumina | 2x Sanger; 14x 454, 20x Illumina | 2.60 | 18.07 | 0.05 | 73.71 |
| mLynCan4_v1.p | *Lynx canadensis* (Canada lynx) | NA | LIC74 | 07/26/19 | PacBio Sequel I; 10X genome; Bionano Genomics; Arima Genomics Hi-C | 72x | 2.41 | 146.11 | 7.50 | 6.18 |
| PanLeo1.0 | *Panthera leo* (lion) | NA | Brooke | 10/07/19 | Illumina; Oxford Nanopore; 10X Genomics | 46x | 2.39 | 136.05 | 0.29 | 242.29 |
| ASM864105v1 | *Canis lupus familiaris* (dog) | German Shepherd | Nala | 09/25/19 | PacBio Sequel; Oxford Nanopore PromethION; Illumina (10X Chromium) | 30x | 2.40 | 64.35 | 20.91 | 22.83 |
| ASM488618v2 | *Canis lupus familiaris* (dog) | Basenji | MU ID 185726 | 08/16/19 | Sequel | 45x | 2.41 | 61.09 | 3.13 | 120.80 |
| UMICH_Zoey_3.1 | *Canis lupus familiaris* (dog) | Great Dane | Zoey | 05/30/19 | PacBio RSII | 50x | 2.34 | 64.20 | 4.72 | 16.82 |
| CanFam3.1 | *Canis lupus familiaris* (dog) | boxer | Tasha | 11/02/11 | Sanger | 7x plus >90Mb finished sequence | 2.39 | 45.88 | 0.27 | 75.10 |

2 <sup>1</sup>All species-specific assembly metrics derived from the NCBI assembly archive.

### Sequence accuracy and quality assessment

Illumina whole-genome sequence data from Cinnamon was used to identify reference sequence errors as homozygous SNVs. These numbered 60,449 in total, indicating a high level of sequence accuracy across assembled contigs (>99.9%). Sequence order and orientation was also highly accurate (>98%), as only 1.2% of BAC-end sequence alignments derived from Cinnamon were identified as discordant. Felis_catus_9.0 sequence order and orientation was also supported by high levels of agreement between individual chromosome sequence alignment and ordered sequence markers from the published cat genetic linkage map [16] (**S1 Data**). The raw sequence data, assembled contigs, and sequence order coordinates (AGP) were accessioned and are fully available by searching GCF_000181335.3 in GenBank.

### Gene annotation

The number of annotated protein-coding genes was very similar between the NCBI and Ensembl pipelines at 19,748 and 19,409, respectively. Approximately 376 protein-coding genes (NCBI) were identified as novel with no matching annotations in Felis_catus*_*8.0 (**S2 Data**). Conversely, 178 genes from Felis_catus*_*8.0 did not map to Felis_catus*_*9.0, of which the cause is unknown (**S1 Table**). A large portion of genes changed substantially (8.4%) in Felis_catus_9.0 during NCBI annotation (**S2 Data**). Aligned sequence of known same-species RefSeq transcripts (n = 420) to Felis_catus*_*9.0 is higher (99.5%) than Felis_catus*_*8.0 (97.8%) and the mean coverage of these same translated RefSeq proteins is also improved (90.1% versus 88.3% in Felis_catus*_*8.0). One important consequence of the less fragmented gene annotation is a 2% increase in aggregate sequence alignments of feline RNA-seq datasets to Felis_catus*_*9.0. These improvements are largely attributed to fewer assembly gaps. The various reported metrics of gene annotation quality conclusively show the protein-coding genes of the domestic cat are of high quality for all trait discovery studies.

### Genetic variation in cats

To improve variant knowledge of the domestic cat, variants from a diverse set of 74 resequenced cats from the 99 Lives project were analyzed in depth. The average sequence coverage was 38.5x with a mean of 98% reads mapped per cat (**S3 Data**). Approximately 46,600,527 variants were discovered, 39,043,080 were SNVs with 93% as biallelic displaying a Ts/Tv ratio of 2.44, suggesting a relatively high level of specificity for variant detection (**S2 Table**). In addition, 97% of SNV positions from the feline 63K genotyping array mapped to Felis_catus_9.0 were detected as SNVs in the WGS call set (**S3 Table**; **S4 Data**) [26]. Using the variant data to estimate cat relatedness, 13 highly related cats (□ > 0.15), two cats with poor read quality, four bengals or bengal crosses, and Cinnamon, the reference, were removed from the sequence dataset to obtain a final set of 54 cats for all subsequent analyses (**S3 Data**). The average number of discovered SNVs per cat was 9.6 million (**Fig. 1a**). Differences in SNV numbers varied according to whether cats were from a specific breed or were random bred (P-value < 0.005, Wilcoxon rank sum test), the two cats with the lowest number of SNVs (∼8 million) were both Abyssinians, the same breed as Cinnamon, while random bred cats from either the Middle East or Madagascar each carried the highest number (> 10.5 million) (**S5 Data**). Individual singleton frequency and estimated inbreeding coefficients (*F* statistic) showed a similar trend with random bred cats generally having significantly more singletons and higher levels of heterozygosity than breed cats (P-value < 0.005, Wilcoxon rank sum test) (**Fig. 1b, Fig. 1c**). Breed cats with higher levels of variation and heterozygosity were either from newly established breeds or were outcrossed individuals. For example, Napoleon cats, which all had an *F* statistic at least one standard deviation below the mean (**Fig. 1c; S5 Data**), are frequently outcrossed as their defining trait dwarfism is likely homozygous lethal *in utero* [27].

**Figure 1.**
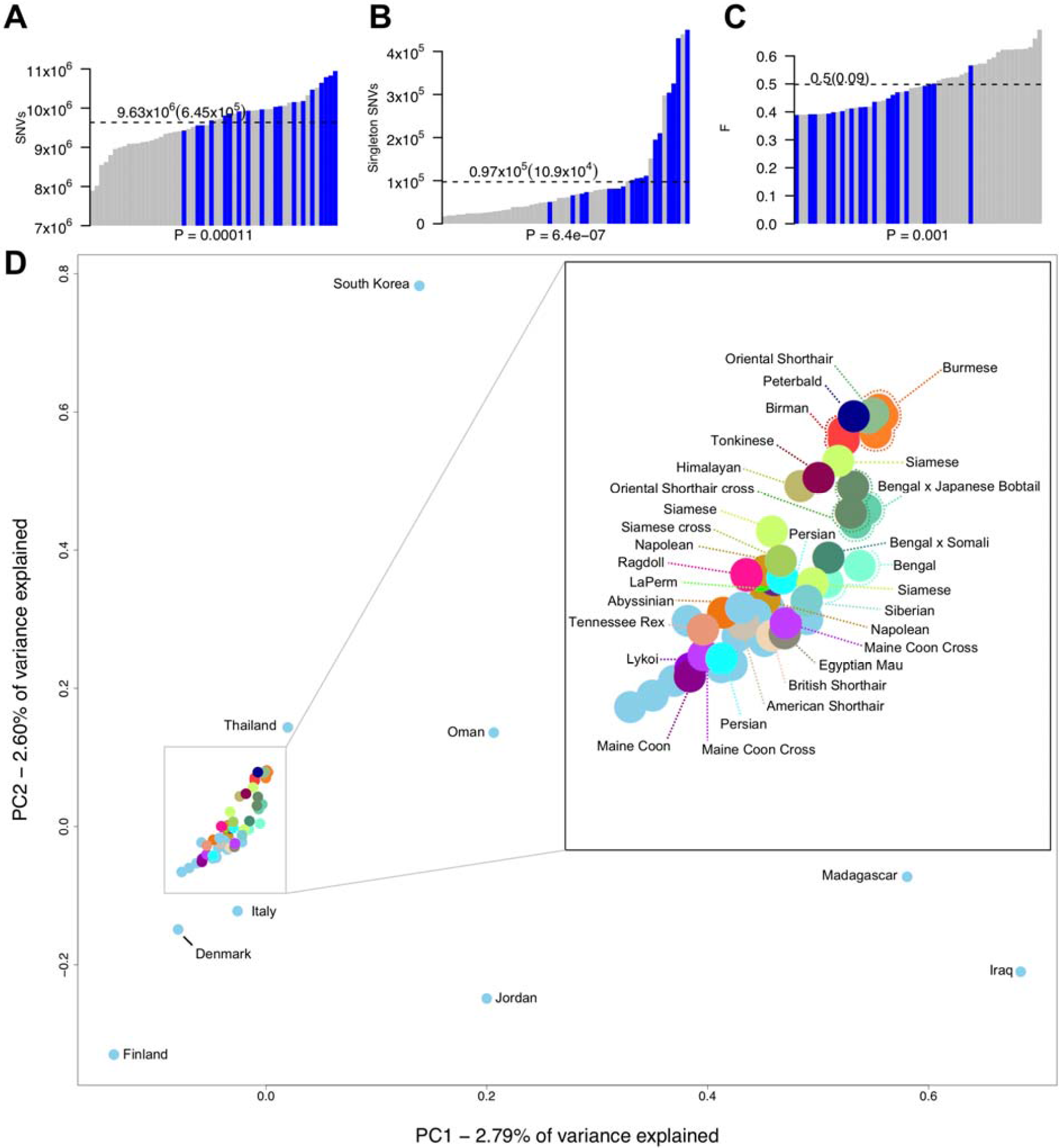
Genetic variation in whole genome sequenced cats. **(a)** Number of SNVs found in each genome, **(b)** singletons per genome, and **(c)** the per sample inbreeding coefficient *F*. Blue bars indicate each individual random bred cat and grey bars indicate each individual breed cat. P-values underneath each x axes were calculated using Wilcoxon rank-sum test and were used to compare breed cats to random bred cats. Dotted lines indicate the mean for each statistic, which is printed above along with standard deviation in braces. **(d)** Population structure of all unrelated cats (including Bengal breeds) estimated using principal components analysis. Random bred cats are colored light blue. Those that were sampled globally for diversity regions are named according to their sampling location. All other cats are named according to their breed or breed cross.

PCA analysis showed the expected distribution of genetic relatedness among cats when considering their geographical location and genetic origins (**Fig. 1d**). In general, most random bred cats displayed a scattered distribution consistent with previous studies on cat population diversity and origins [26, 28]. Although tightly clustered, breed cats could also be distinguished according to their populations of origin. The Asian-derived breeds, Siamese, Burmese, Birman, and Oriental shorthairs were at one end of the spectrum, clustering closely with random bred cats from Thailand. Conversely, cats derived from western populations, such as Maine Coons and Persians, were at the opposite end of the spectrum grouping with random bred cats from Northern Europe, such as Denmark and Finland.

### Implications of feline genetic variation on human disease genes

To characterize feline genetic variation in disease contexts, variant effect predictor (VEP) was used to identify 128,844 synonymous, 77,662 missense, and 1,179 loss of function (LoF) SNVs, where SNVs causing a stop gain were the largest contributor to the LoF category (**S4 Table**). In addition to SNP annotation, genes were grouped according to their genetic constraint across human populations, where genetic constraint was expressed as probability of LoF intolerance. In total, 15,962 cat-human orthologs were identified with 14,291 assigned pLI values. Of these, 9,299 were in the weak constraint group (pLI < 0.1), 2,946 were in the moderate constraint group (0.1 < pLI < 0.9), and 2,739 were in the strong constraint group (pLI > 0.9). For genes under weak constraint in humans, feline SNV density within coding sequences, regardless of impact on gene function, was similar to expected SNV densities based on random assignment of SNV impacts. Conversely, the density of SNVs in genes under strong constraint varied significantly according to SNV impact. LoF and missense SNVs, which have potential to deleteriously impact gene function, were depleted by 59.7% and 35.5%, respectively, while synonymous SNVs, which likely have no deleterious impact on gene function, were enriched by 19.2% relative to expected levels (**Fig. 2a**). Similar results, while less pronounced, were also observed for synonymous and missense SNVs in genes under moderate constraint.

**Figure 2.**
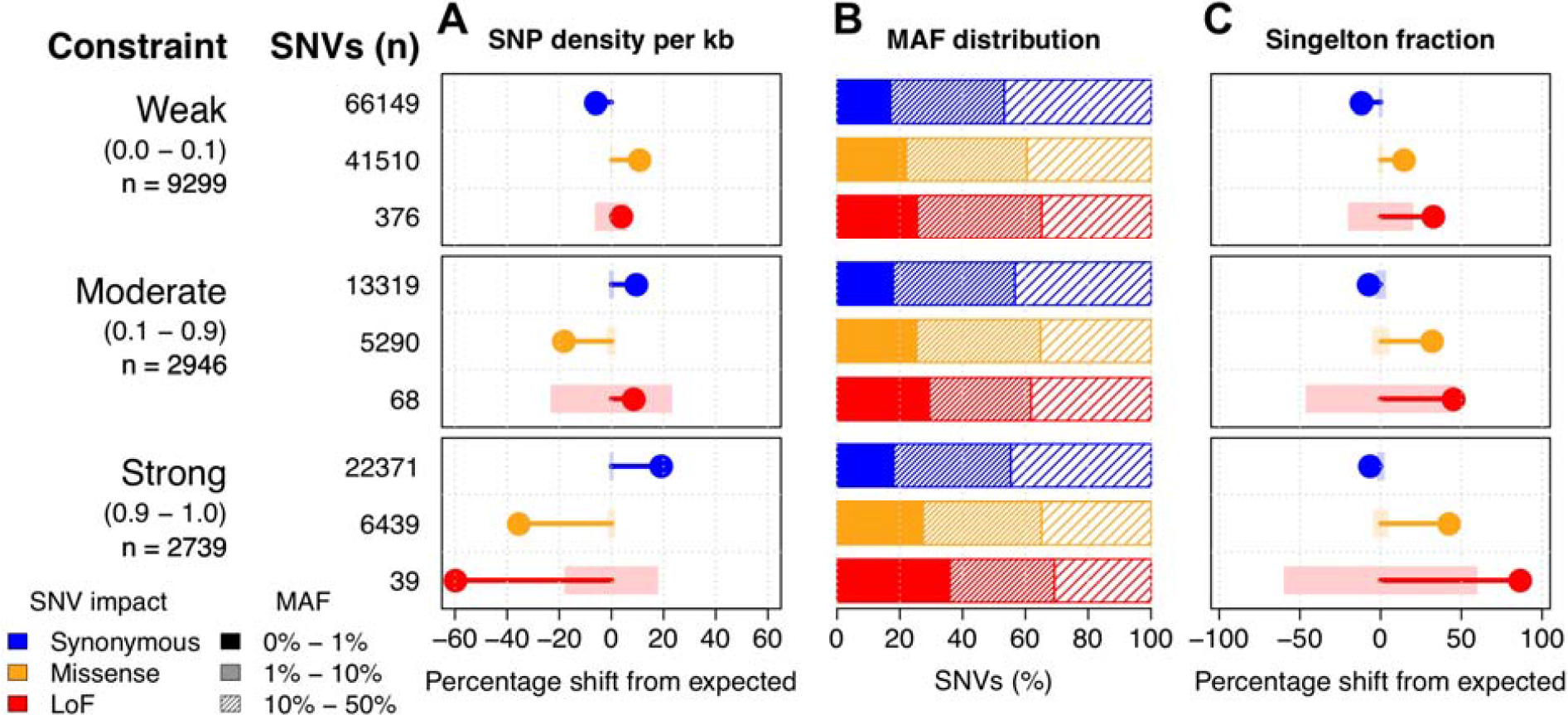
Deleterious SNVs in cats are uncommon and depleted from human constrained genes. To the left of the figure panels, constraint groups are labeled with pLI ranges shown below in braces. Below the pLI ranges are the number of genes found in each constraint group. SNV values show the total number of SNVs of a particular impact that belong to each constraint group. **(a)** Observed percentage differences from expected values for per kb SNV density. Light colored rectangles represent the 95% confidence intervals for the expected values. Confidence intervals were calculated using 10,000 permutations (methods). **(b)** The percentage of SNVs of different impacts and constraint groups across various MAF intervals. **(c)** Observed percentage differences from expected values for the fraction of SNVs in a singleton state (allele count of 1). Light colored rectangles represent the 95% confidence intervals for the expected values, generated from 10,000 permutations.

SNV minor allele frequency (MAF) distributions were compared across each constraint group. MAFs for synonymous SNVs were similarly distributed in all constraint groups. Conversely, the distribution of nonsynonymous SNVs increasingly skewed toward lower MAFs under stronger levels of constraint. For example, 25.5% of LoF SNVs in genes under weak constraint have a MAF < 1%, whereas 35.9% of LoF SNVs in genes under strong constraint have a MAF < 1% (**Fig. 2b**). To determine whether these shifts in MAFs were significant, the fraction of SNVs in a singleton state were compared to expected levels based on random assignment of singleton states. Singleton states of SNVs were significantly enriched for nonsynonymous SNVs with enrichment levels increasing under stronger constraint, indicating many SNVs with functional impacts in genes under constraint are likely rare (**Fig. 2c**). Together, these results show a significant association between selection in human genes and SNV accumulation in cats, suggesting selection pressure within cats and humans is similar across orthologous genes.

Overall, 16 LoF singleton SNVs were identified in intolerant orthologs as potential candidate disease causing variants. Since some cats within 99 lives had recorded disease statuses, these SNVs were assessed for their potential role in cat diseases (**S5 Table**). Of the 16 SNVs, four were supported by both Ensembl and NCBI annotations and were in cats segregating for particular diseases (**Table 2**). Of note is a stop gain in the tumor suppressor *F-box and WD repeat domain containing 7* (*FBXW7*) [29], which was only found in a parent and child segregating for feline mediastinal lymphoma. Other LoF SNVs include stop gains found in *Family With Sequence Similarity 13 Member B* (*FAM13B*) in a random bred with ectodermal dysplasia, cytoplasmic FMR1 interacting protein 2 (*CYFIP2*) in an Egyptian Mau with urate stones, and *SH3 And PX Domains 2A* (*SH3PXD2A*) in a random bred cat with feline infectious peritonitis. Most candidates are not likely disease causing, as each cat carried a mean of 10.0 LoF SNVs in strongly constrained genes (**S1 Fig**.). However, while most LoF SNVs had MAFs > 10%, the mean number of LoF SNVs with MAF < 1% in strongly constrained genes was 0.26 per cat (**S2 Fig**.). These results suggest gene intolerance to mutations may provide as a useful metric for reducing the number of candidate variants for certain diseases.

**Table 2.** High impact singletons in intolerant orthologs for unrelated cats with disease traits.

| SNV location<sup>a</sup> | Ref/Alt | Consequence | Gene symbol | pLI | Individual ID | Disease | Status |
| --- | --- | --- | --- | --- | --- | --- | --- |
| chrA1:116653102 | G/A | Stop gained | *FAM13B* | 0.92 | felCat.Fcat19194.Pudge | Ectodermal dysplasia<sup>b</sup> | Affected |
| chrA1:192824838 | G/A | Stop gained | *CYFIP2* | 1.00 | felCat.Fcat20406.Gannon | Stones | Affected |
| chrB1:77305060 | C/T | Stop gained | *FBXW7* | 1.00 | felCat.Fcat5012.Colorado<sup>c</sup> | Lymphoma | Carrier |
| chrD2:63980875 | G/A | Stop gained | *SH3PXD2A* | 1.00 | felCat.CR1397.Isabella | Infectious peritonitis<sup>d</sup> | Affected |

<sup>a</sup> SNV locations were only reported if they were supported by NCBI annotations.
<sup>b</sup> A second unrelated cat, felCat.Fcat19197.Kooki, was also affected.
<sup>c</sup> Affected offspring removed earlier from analysis inherited the same SNV.
<sup>d</sup> A second unrelated cat, felCat.CR1219.Tamborine, was also affected.

### Structural variant discovery

The merging of the two independent SV call sets was performed across all individuals for variants occurring within 50 bp of the independent variant call position, with agreement on variant type and strand, and variant size within 500 bp. Per cat, an average of 44,990 SVs were identified, with variants encompassing 134.3 Mb across all individuals. Deletions averaged 905 bp, duplications 7,497 bp, insertions 30 bp, and inversions 10,993 bp. The breed and breed crosses (n = 36) compared to random bred cats (n = 18) showed comparable SV diversity (t-test p = 0.6) (**S3 Fig**.). In total, 208,135 SVs were discovered, of which 123,731 (60%) were deletions (**Fig. 3a**). SV population frequencies were similar across SV types, except for inversions. For deletions, duplications and insertions, 38% to 48% of each SV type was found at population frequencies of 0.02 – 0.10 and 0.10 – 0.50. Meanwhile, > 90% of inversions are found at a population frequency of 0.02 – 0.10 (**Fig. 3a**). The majority of SVs identified are common across cats, suggesting their impacts are mostly tolerated.

**Figure 3.**
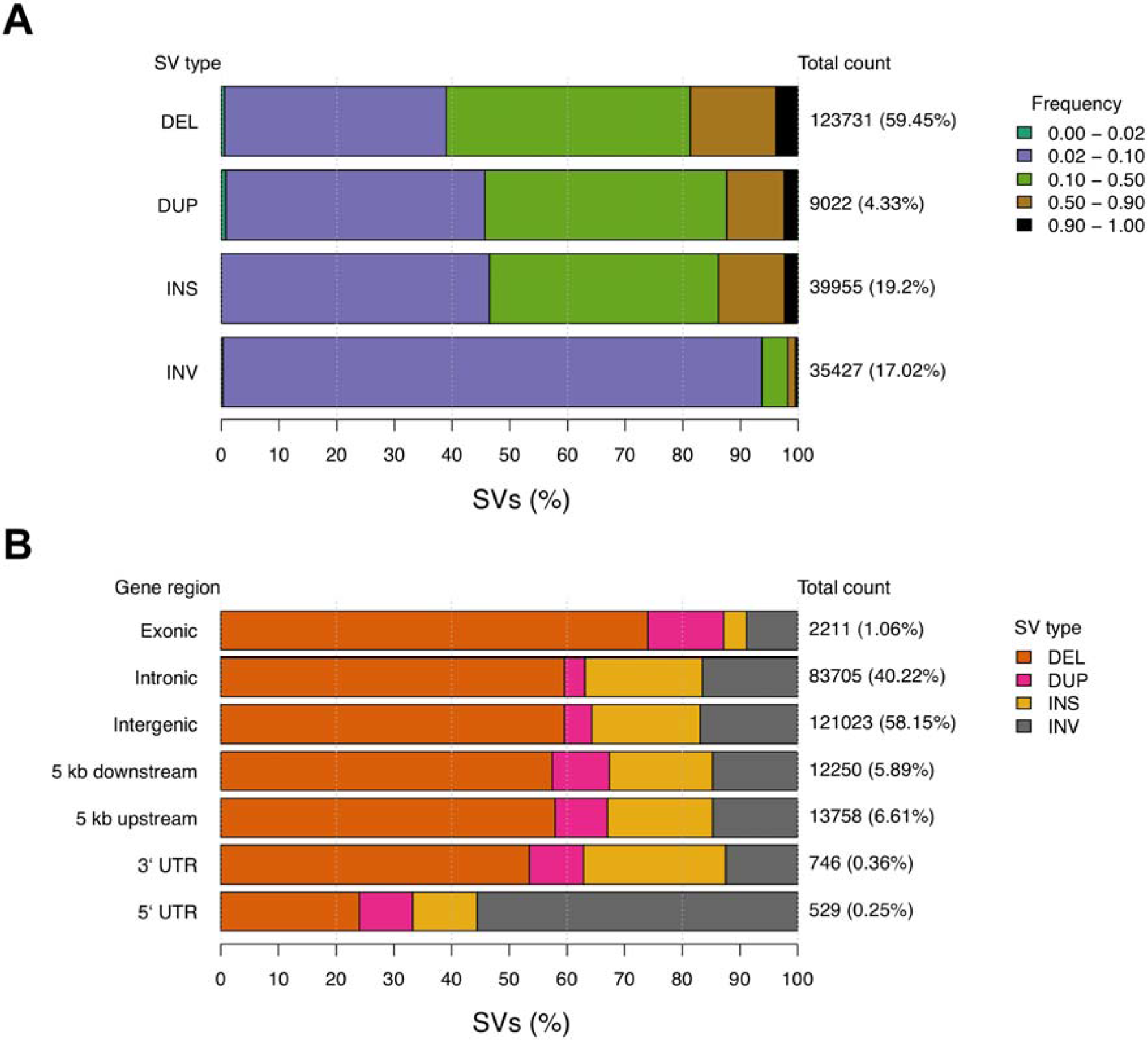
The structural variant landscape of cats. **(a)** Population frequency of each SV type. Colored bars represent the proportion of a given SV type found within various population frequency ranges. Total count values to the right of each bar represent the total number of SVs of each type, while percentage values in braces represent the proportion of all SVs that belong to each type. Frequency ranges shown in the legend are ordered along each bar from left to right. **(b)** The proportion of SVs found in each genomic region. Colored bars represent the proportion of SVs belonging to a particular type found across different genomic regions. Similar to above, the total count values to the right of each bar represent the total number of SVs found in each genomic region. The percentage values in braces represent the genome-wide proportion of all SVs found in each gene region. These values sum greater than 100 percent as a single SV can span multiple types of genomic regions. Structural variants are noted as: deletion (DEL); duplication (DUP); insertion (INS); inversion (INV).

SV density across autosomes was relatively constant with chromosome E1 carrying the largest SV burden at 96.95 SVs per Mb (**S4 Fig**.). Approximately 6,096 SVs (3%, 10.1 Mb) were observed in >90% of the cat genomes (**S6 Data**), indicating the cat used for the Felis_catus_9.0 assembly, Cinnamon, an Abyssinian, likely carries a minor allele at these positions. SV annotation showed that SV counts per region were consistent with the fraction of the genome occupied by each region type. For example, 58.15% of SVs were intergenic, 40.22% of SVs were intronic, and 1.06% of SVs were exonic, potentially impacting 217 different protein coding genes (**Fig. 3b**) (**S7 Data**). Conversely, the proportion of some SV types found in certain gene regions varied from their genome-wide averages. For example, in regions 5 kb upstream and downstream of genes, duplications were increased approximately two-fold. For exonic regions, 74% of SVs were deletions, an increase form the genome wide level of 59.45%. For 5` UTRs, the majority of SVs were inversions, which only represent 17.02% of total SVs. These results suggest an interaction between the impact of SV types and the potential function of the gene regions they are found in.

### Genetics of feline dwarfism

Dwarfism in cats is the defining feature of the munchkin breed and is characterized by shortened limbs and normal sized torso (**Fig. 4a**) [30]. Similar to analyses with cat assemblies Felis_catus-6.2 and Felis_catus_8.0, previous investigations for SNVs in the new Felis_catus_9.0 assembly did not identify any high priority candidate variants for disproportionate dwarfism [27]. However, SV analysis within the critical region previously identified by linkage and GWAS on chromosome B1:170,786,914-175,975,857 [27] revealed a 3.3 kb deletion at position chrB1:174,882,897-174,886,198, overlapping the final exon of *UDP-glucose 6-dehydrogenase (UGDH)* (**Fig. 4b**). Upon manual inspection of this SV, a 49 bp segment from exon 8 appeared to be duplicated and inserted 3.5 kb downstream, replacing the deleted sequence. This potentially duplicated segment was flanked by a 37 bp sequence at the 5` end and a 20 bp sequence at the 3’ end, both of unknown origin (**Fig. 4c**). Discordant reads consistent with the SV were private to all three unrelated WGS dwarf samples (**S5 Fig**.). The breakpoints surrounding the deletion were validated in WGS affected cats with Sanger sequencing. PCR-based genotyping of the 3.3 kb deletion breakpoints was conducted in an additional 108 cats including, 40 normal and 68 affected dwarf cats (**S6 Fig**.). Expected amplicon sizes and phenotypes were concordant across all cats, except for a “munchkin; non-standard (normal legs); Selkirk mix”, which appeared to carry the mutant allele, suggesting an alternate causal gene or sampling error.

**Figure 4.**
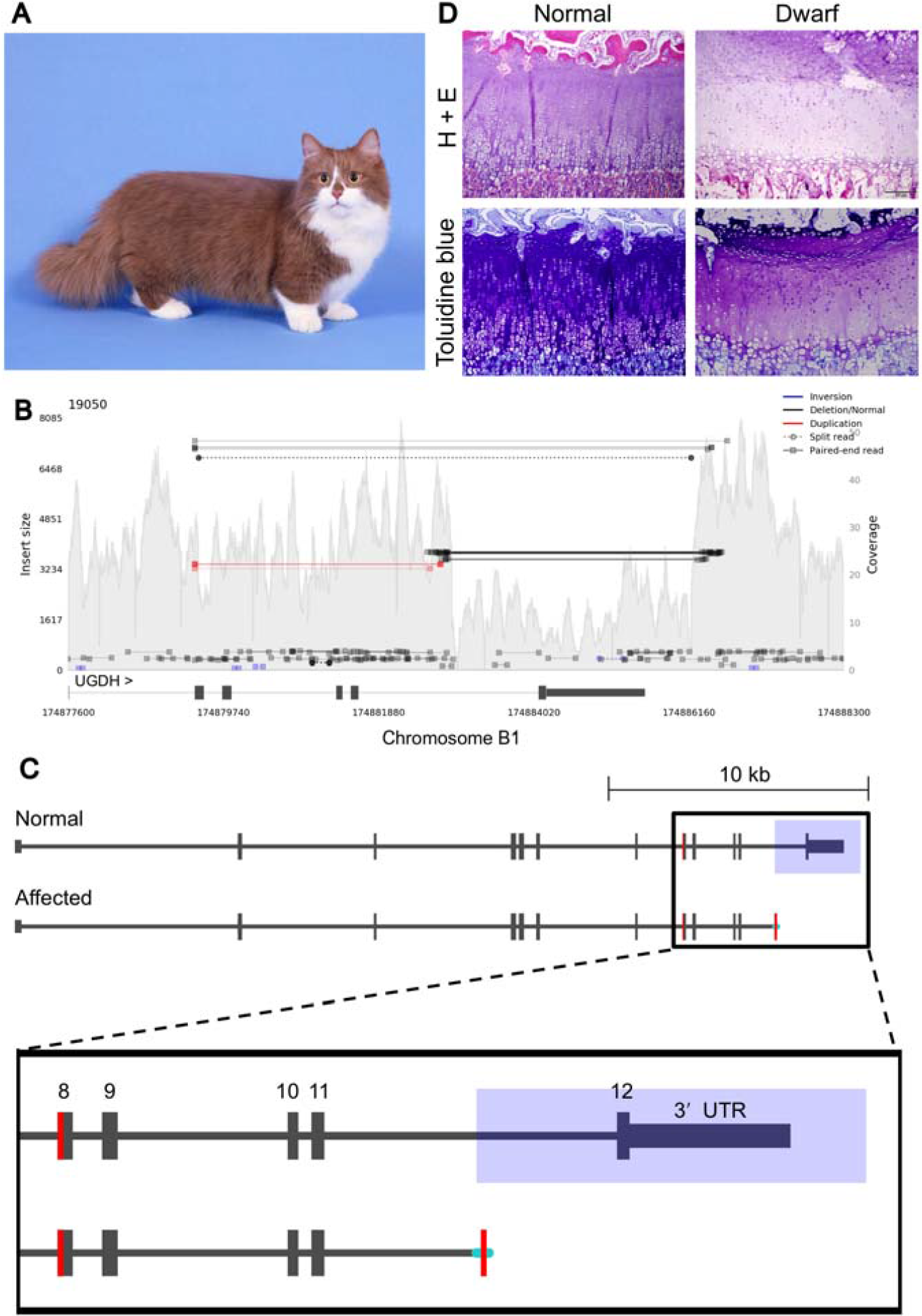
A complex structural variant is associated with feline dwarfism. **(a)** Image of dwarf cat from the munchkin breed. Notice that limbs are short while torso remains normal size. Image was donated courtesy of Terri Harris. **(b)** Samplot output of discordant reads within an affected cat. Decreased coverage over the final exon of *UGDH* represents a deletion. Discordant reads spanning beyond the deleted region show newly inserted sequence sharing homology with *UGDH* exon 8. **(c)** Schematic illustrating the candidate structural variant for dwarfism. Blue rectangle overlaps deleted region, shaded portion of exon 8 shows potential duplication in the affected, and cyan lines indicate inserted sequence of unknown origin. **(d)** Hematoxylin and eosin and toluidine blue histologic samples of the distal radius epiphyseal cartilage plate from a neonatal kitten and an age-matched dwarf kitten. The normal kitten showed regular columnar arrangement of chondrocytes with abundant proteoglycan as determined by strong metachromasia with toluidine blue stain. The dwarf kitten showed disorganized columnar arrangement of chondrocytes with proteoglycan depletion as determined by weak metachromasia with toluidine blue stain.

As *UGDH* has known roles in proteoglycan synthesis in chondrocytes [31, 32], growth plates in two healthy and seven dwarf neonatal kittens were histologically analyzed for structural irregularities and proteoglycan concentrations (Methods). The articular cartilage or bone in the dwarf specimens had no significant histopathologic changes and epiphyseal plate was present in all specimens. In the two normal kittens, chondrocytes in the epiphyseal plate exhibited a regular columnar arrangement with organization into a zone of reserve cells, a zone of proliferation, a zone of hypertrophy and a zone of provisional calcification (**Fig. 4d; S7 Fig**.). Moreover, the articular-epiphyseal cartilage complex and the epiphyseal plate in the distal radius from these two normal kittens also contained abundant proteoglycan as determined by toluidine blue stain. In contrast, in dwarf kittens, the epiphyseal plate had a disorganized columnar arrangement that was frequently coupled with proteoglycan depletion (**Fig. 4d; S7 Fig**.).

## Discussion

Studies of the domestic cat show the amazing features of this obligate carnivore, including their incomplete domestication, wide range of coat color variation, and use as biomedical models [11, 33-35]. Using a combination of long-reads, combined with long-range scaffolding methods, Felis_catus_9.0 was generated as a new cat genome resource, with an N50 contig length (42 Mb) surpassing all other carnivore genome assemblies. This sequence contiguity represents a 1000-fold increase of ungapped sequence (contigs) length over Felis_catus*_*8.0, as well as a 40% reduction in the amount of unplaced sequence. All measures of sequence base and order accuracy, such as the low number of discordant BAC end sequence alignments, suggest this reference will strongly support future resequencing studies in cats. Equally important are the observed improvements in gene annotation including the annotation of new genes, overall more complete gene models and improvements to transcript mappability. Only 178 genes were missing in Felis_catus*_*9.0 compared to Felis_catus_8.0, which will require further investigation. However, 376 predicted genes are novel to Felis_catus_9.0, i.e. not found in Felis_catus_8.0.

Using this Felis_catus_9.0 assembly, a vast new repertoire of SNVs and SVs were discovered for the domestic cat. The total number of variants discovered across our diverse collection of domestic cats was substantially higher than previous studies in other mammals, such as, cow [36], dog [37], rat [38-40], sheep [41, 42], pig [43], and horse [44]. Even rhesus macaques, with twice as many variants as human, do not approach the same levels of cat SNV variation [45, 46]. The cats had 36.6 million biallelic SNVs with individuals carrying ∼9.6 million SNVs each. Conversely, humans roughly carry 4 to 5 million SNVs per individual [47]. One possible explanation for this discrepancy may be the unique process cats underwent for domestication. Rather than undergoing strong selective breeding leading to a severe population bottleneck, cats were instead “self” domesticated, never losing some ancestral traits such as hunting behavior [48-51]. The practice of strong selective breeding in cats only began in recent history and is based almost exclusively on aesthetic traits. Consistent with previous analyses, our evaluation of breeds and random bred cats showed tight clustering between cats from different breeds that suggest cat breeds were likely initiated from local random bred cat populations [28, 52]. In regards to per sample numbers of variants, most breed cats had fewer SNVs than random bred cats. Likewise, random bred cats also had higher numbers of singletons and a lower inbreeding coefficient than breed cats, suggesting breed cats may share a common genetic signature distinct from random bred populations.

For genetic discovery applications, animals with lower levels of genetic diversity are often more desirable as they reduce experimental variability linked to variation in genetic architecture [53]. However, with decreasing costs of genome or exome sequencing leading to exponential growth in variant discovery, animals with higher genetic diversity, such as cats, could enhance discovery of tolerable loss of function or pathogenic missense variants. Similar to cats, rhesus macaques from research colonies, exhibited per sample SNV rates more than two-fold greater than human [46]. Importantly, macaque genetic diversity has been useful for characterizing non-deleterious missense variation in humans [54].

A major premise of our feline genomics analysis is that increased discovery of genetic variants in cats will help improve benign/pathogenic variant classification and reveal similarities between human and feline genetic disease phenotypes, aiding disease interpretation in both species. To gain insights into the burden of segregating variants with potentially deleterious impacts on protein function, SNV impacts were classified across 54 unrelated domestic cats. Overall, cat genes identified as being under strong constraint in humans were depleted of nonsynonymous SNVs and enriched for potentially rare variants, showing both species harbor similar landscapes of genetic constraint and indicating the utility of Felis_catus_9.0 in modeling human genetic disease. An important limitation on the analysis was the available sample size. As there were only 54 cats, true rare variants, defined as variants with allele frequency of at least 1% in the population [55], could not be distinguished from common variants that appear in only one of 54 randomly sampled cats by chance. Instead, singleton SNVs were focused on as candidates for rare variants. Altogether, 18.6% of all feline SNVs could be considered candidates for rare variants. Alternatively, larger analyses in human reveal a much higher fraction of low frequency variants [56, 57]. For example, 36% of variants in an Icelandic population of over 2000 individuals had a minor allele frequency below 0.1% [57]. Similarly sized analyses in other mammals reported ∼20% of variants had allele frequencies < 1% [42, 44, 46]. As the number of cats sequenced increases, the resolution to detect rare variants, as well as the total fraction of low frequency variants, will continue to grow linearly as each individual will contribute a small number of previously undiscovered variants [58].

Focusing specifically on potentially rare variants with high impacts in constrained genes identified a potential cause for early onset feline mediastinal lymphoma, a stop gain in tumor suppressor gene *FBXW7* [29]. Feline mediastinal lymphoma is distinct from other feline lymphomas in its early onset and prevalence in Siamese cats and Oriental Shorthairs [59], suggesting a genetic cause for lymphoma susceptibility specific to Siamese related cat breeds [60, 61]. The stop gain was initially observed in a heterozygous state in a single cat identified as a carrier in the unrelated set of cats. Subsequent analysis of the full set of cats revealed the SNV had been inherited in an affected offspring. Despite discordance between the presence of this mutant allele and the affection status of the cats it was found in, the variant still fits the profile as a susceptibility allele for mediastinal lymphoma. For example, homozygous knockout of *Fbxw7* is embryonic lethal in mice [62, 63], while heterozygous knockout mice develop normally [63]. However, irradiation experiments of *Fbxw7*^+/−^ mice and *Fbxw7*^+/-^*p53*^+/-^ crosses identify *Fbwx7* as a haploinsufficient tumor suppressor gene that requires mutations in other cancer related genes for tumorgenesis [64], a finding supported in subsequent mouse studies [65, 66]. Similarly, in humans, germline variants in *FBXW7* are strongly associated with predisposition to early onset cancers, such as Wilms tumors and Hodgkin’s lymphoma [67, 68]. Screening of Siamese cats and other related breeds will validate the *FBWX7* stop gain as a causative mutation for lymphoma susceptibility and may eventually aid in the development of a feline cancer model.

An important concern with using human constraint metrics for identifying causative variants in cats is the potential for false positives. For example, out of 16 SNVs initially identified as potential feline disease candidates, many belonged to cats with no recorded disease status (**S5 Table**). However, since a large fraction of healthy humans also carried SNVs matching similar disease causing criteria as used in cats [56], evolutionary distance between humans and cats is unlikely to be a significant contributing factor to false positive candidate disease variant identification. Instead, the frequency of high impact variants in constrained genes is likely due to the limited resolution provided by analyses at the gene level. Recent strategies in humans have confronted this problem by focusing on constraint at the level of gene region. These analyses identified many genes with low pLI values that contained highly constrained gene regions that were also enriched for disease causing variants [69]. Alternatively, another strategy for further refining constrained regions could involve combining genomic variation from multiple species. Many variants observed in cats, along with their impact on genes, are likely unique to cats and could potentially be applied to variant prioritization workflows in humans.

Presented here is the first comprehensive genome-wide SV analysis for Felis_catus_9.0. Previous genome-wide SV analyses in the cat were performed in Felis_catus-6.2 and focused solely on copy number variation (CNV) [24]. Approximately 39,955 insertions, 123,731 deletions, 35,427 inversions, and 9,022 duplications were identified, far exceeding previous CNV calculations of 521 losses and 68 gains. This large discrepancy is likely due to a number of reasons including filtering stringency, sensitivity, and reference contiguity, where increased numbers of assembly gaps in previous reference genome assemblies hindered detection of larger SV events. An important difference regarding filtering stringency and sensitivity is that the average CNV length was 37.4 kb as compared to 2.7 kb for SVs in Felis_catus_9.0. Since CNV detection depends on read-depth [70, 71], only larger CNVs can be accurately identified, as coverage at small window sizes can be highly variable. The use of both split-read and read-pair information [72, 73], allowed the identification of SV events at much finer resolution than read-depth based tools [74].

Improved sensitivity for smaller SV events, while helpful for finding new disease causing mutations, may also lead to increased false positive SV detection. In three human trios sequenced with Illumina technology, LUMPY and Delly were both used to identify 12,067 and 5,307 SVs respectively, contributing to a unified call set, along with several other tools, of 10,884 SVs per human on average [75]. In cats, the average number of SVs per individual was 4 times higher than in humans, with 44,990 SVs per cat, suggesting the total number SVs in cats is likely inflated. However, the majority of the SVs were at population frequencies below 0.5, ruling out poor reference assembly as a contributing factor. Instead, the difference in SV count between cats and humans is likely due to sample specific factors. For example, the majority of samples were sequenced using two separate libraries of 350 bp and 550 bp insert sizes (**S2 Data**). Despite the potentially high number of false positive SVs, increased SV sensitivity was useful for trait discovery. The deletion associated with dwarfism was only found in the Delly2 call set. If SV filtering were more stringent, such as requiring SVs to be called by both callers, the feline dwarfism SV may not have otherwise been detected. Ultimately, these results highlight the importance of high sensitivity for initial SV discovery and the use of highly specific molecular techniques for downstream validation of candidate causative SVs.

The improved contiguity of Felis_catus*_*9.0 was particularly beneficial for identifying a causative SV for feline dwarfism. In humans, approximately 70% of cases are caused by spontaneous mutations in *fibroblast growth factor 3* resulting in achondroplasia or a milder form of the condition known as hypochondroplasia [76, 77]. The domestic cat is one of the few species with an autosomal dominant mode of inheritance for dwarfism that does not have other syndromic features. It can therefore provide as a strong model for hypochondroplasia. Previous GWAS and linkage analyses suggested a critical region of association for feline disproportionate dwarfism on cat chromosome B1 that spanned 5.2 Mb [27]. Within the critical region, SV analysis identified a 3.3 kb deletion that had removed the final exon of *UGDH*, which was replaced by a 106 bp insertion with partial homology to *UGDH* exon 8, suggesting a potential duplication event. Importantly, *UGDH* likely plays a role in proteoglycan synthesis in articular chondrocytes, as osteoarthritic human and rat cartilage samples have revealed reduced UGDH protein expression was associated with disease state [31]. Similarly, in dwarf cat samples, histology of the distal radius showed irregularity of the chondrocyte organization and proteoglycan depletion. Collectively, results suggest a disease model of reduced proteoglycan synthesis in dwarf cat chondrocytes caused by loss of function of *UGDH* resulting in abnormal growth in the long bones of dwarf cats. In ClinVar, only one missense variant in *UGDH* has been documented <u>NM_003359.4(</u>**<u>UGDH</u>**<u>):c.950G>A (p.Arg317Gln)</u> and is associated with epileptic encephalopathy. Seventeen other variants involving *UGDH* span multiple genes in the region. The SV in the dwarf cats suggest *UGDH* is important in early long bone development. As a novel gene association with dwarfism, *UGDH* should be screened for variants in undiagnosed human dwarf patients.

High-quality genomes are a prerequisite for unhindered computational experimentation. Felis_catus_9.0 is currently the most contiguous genome of a companion animal, with high accuracy and improved gene annotation that serves as a reference point for the discovery of genetic variation associated with many traits. This new genomic resource will provide a foundation for the future practice of genomic medicine in cats and for comparative analyses with other species.

## Methods

### Whole genome sequencing

The same genome reference inbred domestic cat, Cinnamon, the Abyssinian, was used for the long-read sequencing [78] [33]. High molecular weight DNA was isolated using a MagAttract HMW-DNA Kit (Qiagen, Germantown, MD) from cultured fibroblast cells according to the manufacturer’s protocol. Single molecule real-time (SMRT) sequencing was completed on the RSII and Sequel instruments (Pacific Biosciences, Menlo Park, CA).

### Genome assembly

All sequences (∼72x total sequence coverage) were assembled with the fuzzy Bruijn graph algorithm, WTDBG (https://github.com/ruanjue/wtdbg), followed by collective raw read alignment using MINIMAP to the error-prone primary contigs [79]. As a result, contig coverage and graph topology were used to detect base errors that deviated from the majority haplotype branches that are due to long-read error (i.e. chimeric reads) or erroneous graph trajectories (i.e. repeats) as opposed to allelic structural variation, in which case, the sequence of one of the alleles is incorporated into the final consensus bases. As a final step to improve the consensus base quality of the assembly, from the same source DNA (Cinammon), short read sequences (150 bp) were generated from 400 bp fragment size TruSeq libraries to ∼60X coverage on the Illumina HiSeqX instrument, which was then used to correct homozygous insertion, deletion and single base differences using PILON [80].

### Assembly scaffolding

To generate the first iteration of scaffolds from assembled contigs, the BioNano Irys technology was used to define order and orientation, as well as, detect chimeric contigs for automated breaks [81]. HMW-DNA in agar plugs was prepared from the same cultured fibroblast cell line using the BioNano recommended protocol for soft tissues, where using the IrysPrep Reagent Kit, a series of enzymatic reactions lysed cells, degraded protein and RNA, and added fluorescent labels to nicked sites. The nicked DNA fragments were labeled with ALEXA Fluor 546 dye and the DNA molecules were counter-stained with YOYO-1 dye. After which, the labeled DNA fragments were electrophoretically elongated and sized on a single IrysChip, with subsequent imaging and data processing to determine the size of each DNA fragment. Finally, a *de novo* assembly was performed by using all labeled fragments >150 kb to construct a whole-genome optical map with defined overlap patterns. Individual maps were clustered, scored for pairwise similarity, and Euclidian distance matrices were built. Manual refinements were then performed as previously described [81].

### Assembly QC

The scaffolded assembly was aligned to the latest cat linkage map [16] to detect incorrect linkage between and within scaffolds, as well as discontinuous translocation events that suggest contig chimerism. Following a genome-wide review of interchromosomal scaffold discrepancies with the linkage map, the sequence breakpoints were manually determined and the incorrect sequence linkages were separated. Also, to assess the assembly of expanded heterozygous loci ‘insertions’, the same reference DNA sequences (Illumina short read inserts 300 bp) were aligned to the chromosomes to detect homozygous deletions in the read alignments using Manta [82], a structural variant detection algorithm. In addition, the contigs sequences were aligned to the Felis_catus_8.0 using BLAT [83] at 99% identity and scored alignment insertion length at 0.5 to 50 kb length to further refine putatively falsely assembled heterozygous loci that when intersected with repeat tracks suggested either error in the assembly or inability to correctly delineate the repetitive copy.

### Chromosome builds

Upon correction and completion of the scaffold assembly, the genetic linkage map [16] was used to first order and orient all possible scaffolds by using the Chromonomer tool similarly to the previously reported default assembly parameter settings [84]. A final manual breakage of any remaining incorrect scaffold structure was made considering various alignment discordance metrics, including to the prior reference Felis_catus_8.0 that defined unexpected interchromosomal translocations and lastly paired end size discordance using alignments of BAC end sequences from the Cinnamon DNA source (http://ampliconexpress.com/bac-libraries/ite).

### Gene Annotation

The Felis_catus*_*9.0 assembly was annotated using previously described NCBI [85, 86] and Ensembl [87] pipelines, that included masking of repeats prior to *ab initio* gene predictions and evidence-supported gene model building using RNA sequencing data [88]. RNA sequencing data of varied tissue types (https://www.ncbi.nlm.nih.gov/genome/annotation_euk/Felis_catus/104) was used to further improve gene model accuracy by alignment to nascent gene models that are necessary to delineate boundaries of untranslated regions as well as to identify genes not found through interspecific similarity evidence from other species.

### Characterizing feline sequence Variation

Seventy-four cat WGSs from the 99 Lives Cat Genome Project were downloaded from the NCBI short read archive (SRA) affiliated with NCBI biosample and bioproject numbers (**S2 Data**). All sequences were produced with Illumina technology, on either an Illumina HiSeq 2500 or XTen instrument using PCR-free libraries with insert lengths ranging from 350 bp to 550 bp, producing 100 – 150 bp paired-end reads. WGS data was processed using the Genome analysis toolkit (GATK) version 3.8 [89, 90]. BWA-MEM from Burrows-Wheeler Aligner version 0.7.17 was used to map reads to Felis_catus_9.0 (GCF_000181335.3) [91]. Piccard tools version 2.1.1 (http://broadinstitute.github.io/picard/) was used to mark duplicate reads, and samtools version 1.7 [92] was used to sort, merge and index reads. Tools used from GATK 3.8 consisted of IndelRealigner and RealignerTargetCreator for indel realignment, BaseRecalibrator for base quality score recalibration (BQSR) [93], and HaplotypeCaller and GenotypeGVCFs for genotyping [94]. The variant database used for BQSR was built by first genotyping non-recalibrated BAMs and applying a strict set of filters to isolate high confidence variants. To determine the final variant call set, post BQSR, a less-strict GATK recommended set of filters, was used. All filtering options are outlined in supplementary material (**S6 Table**). The set of unrelated cats was determined using vcftools’ relatedness2 function on SNP genotypes to estimate the kinship coefficient, □, for each pair of cats [95]. First, related cats were identified as sharing potential sibling and parent-child relationships if □ > 0.15. Next, cats with the highest number of relatives were removed in an iterative fashion until no relationships with □ > 0.15 remained. To detect population structure among the sequenced cats, a principal component analyses was conducted using SNPRelate version 1.16.0 in the R Statistical Software package. The SNV set generated after the appropriate quality control measures were used was further filtered for non-biallelic SNVs and sites displaying linkage disequilibrium (r2 threshold = 0.2) as implemented in the SNPRelate package.

### Measuring coding variant impacts on human disease genes

VCF summary statistics and allele counts were computed using various functions from vcftools [95] and vcflib (https://github.com/vcflib/vcflib). SNVs impacts of synonymous, missense, and LoF were determined using Ensembl’s variant effect predictor (VEP) with annotations from Ensembl release 98 [87, 96]. Cat and human orthologs were identified using reciprocal best hits blast, where Ensembl 98 protein fasta sequences were used as queries. Using pLI for human genes obtained from gnomAD [56, 97], genes were assigned to constraint groups based on weak constraint (pLI < 0.1), moderate constraint (pLI > 0.1 and pLI < 0.9), and strong constraint (pLI > 0.9). The observed per kb SNV density, *Y*^*D*^, for each constraint group and SNV impact was calculated as, 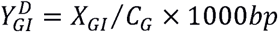, where *C* is the total length of the coding sequence and *X* is the number of SNVs within *C*. The subscript *I* refers to SNV impacts from the subset of all SNVs, *X*. The subscript *G* refers to the subset of either *C* or *X* that are found within a particular constraint group. For example, when *G* represents genes under weak constraint and *I* represents LoF SNVs, *XGI* would be all LoF SNVs within the coding sequence of genes under weak constraint. The expected per kb SNV density, *E*^*D*^, for each constraint group and impact was calculated as, 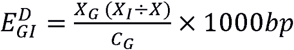. The 95% confidence intervals surrounding the expected per kb SNV densities were calculated from a random distribution generated by 10,000 permutations, where SNV impacts were shuffled randomly across variant sites. The observed singleton fraction, *Y*^*F*^, for each constraint group and impact was calculated as, 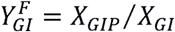, where the subscript *P* refers to SNVs identified as singletons. The expected singleton fraction was calculated as, 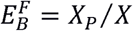, where the 95% confidence intervals surrounding the expected singleton fractions were also calculated from a random distribution generated by 10,000 permutations. For each permutation, SNV MAFs were shuffled so variants were randomly assigned “singleton” status.

### Structural variant identification and analysis

To discover SVs in the size range of <100 kb, aligned reads from all cats to Felis_catus_9.0 were used as input for the LUMPY [73] and Delly2 [72] SV callers. For LUMPY, the empirical insert size was determined using samtools and pairend_distro.py for each BAM. Discordant and split-reads extracted from paired-end data using SpeedSeq [98] were used as input along with each aligned BAM, minimum mapping threshold of 20, and empirical mean and standard deviation of insert size. Samples were called independently. SVTyper [98] was used to genotype each SV before merging all resulting VCFs using BCFtools [92]. For Delly2 [72], all SVs were called for individual BAMs independently, then merged into a single BCF. For each sample, variants were then re-called using the merged results from all samples. The individual re-called VCFs were then merged into a single file using BCFtools. Given the poor resolution of LUMPY for small insertions, only the Delly2 calls were considered and variants were required to be found in more than 2 individuals. A convergence of calls was determined by reciprocal overlap of 50% of the defined breakpoint in each caller as our final set. SVs were annotated using SnpEff [99], which was used to count gene region intersects. SVs were considered exonic if they were annotated as exon_region, frameshift, start_lost, or stop_gained.

### Disease variant discovery for dwarfism

To discover causal variants associated with dwarfism, three unrelated affected cats with disproportionate dwarfism from the 99 Lives genome dataset were examined for SVs. Identified SVs were considered causal candidates if they were, 1) concordant with affection status and an autosomal dominant inheritance pattern, 2) and located within the ∼5.2 Mb dwarfism critical region located on cat chromosome B1:170,786,914 – 175,975,857 [27]. After initial identification, candidate variants were prioritized according to their predicted impact on protein coding genes. High priority candidate SVs were further characterized manually in affected individuals using the integrated genomics viewer (IGV) [100]. STIX (structural variant index) was used to validate candidate SVs by searching BAM files for discordant read-pairs that overlapped candidate SV regions (https://github.com/ryanlayer/stix). After manual characterization, SV breakpoints were validated with PCR amplification and Sanger sequencing. For further genotyping of candidate SVs, all sample collection and cat studies were conducted in accordance with an approved University of California, Davis Institutional Animal Care and Use protocols 11977, 15117, and 16691 and University of Missouri protocols 7808 and 8292. DNA samples from dwarf and normal cats were genotyped for the candidate SV identified in the three sequenced dwarfism cats [27]. PCR primers were designed using the known SV sequence breakpoints (**S6a File**). For validation, PCR amplification products were sanger sequenced and compared against Felis_catus_9.0. For screening, all samples from previous linkage and GWAS studies [27] were genotyped using the primers, UGDH_mid_F, UGDH_del_R, and UGDH_dn_R (**S7 Table**) in a single reaction. PCR products were separated by gel electrophoresis (80V, 90 minutes) in 1.25% (w/v) agarose in 1X TAE. A 622 bp amplicon was expected from the normal allele and a 481 bp amplicon from the affected allele (**S6 Fig**.).

### Histological characterization of dwarf cat growth plates

Cat owners voluntarily submitted cadavers of stillborn dwarf and musculoskeletally normal kittens (7 and 2 respectively, dying from natural causes) via overnight shipment on ice. Distal radius including physis (epiphyseal plate) were collected and fixed in 10% neutral buffered formalin. These tissues were decalcified in 10% EDTA solution. After complete decalcification the tissues were dehydrated with gradually increasing concentrations of ethanol and embedded in paraffin. Frontal sections of distal radial tissues were cut to 6 µm and mounted onto microscope slides. The samples were then dewaxed, rehydrated, and stained with hematoxylin and eosin to evaluate the tissue structure and cell morphology. The sections were also stained with toluidine blue to determine the distribution and quantity of proteoglycans. These samples were subjectively assessed for chondrocyte and tissue morphology and growth plate architecture by a pathologist (KK) who was blinded to the sample information.

## Declarations

### Ethics approval and consent to participate

All sample collection and cat studies were conducted in accordance with an approved University of California, Davis Institutional Animal Care and Use protocols 11977, 15117, and 16691 and University of Missouri protocols 7808 and 8292.

### Availability of data and materials

The datasets supporting the conclusions of this article are available in NCBI’s sequence read archive. The whole genome sequence data generated in this study have been submitted to the NCBI BioProject database (http://www.ncbi.nlm.nih.gov/bioproject/) under accession number PRJNA16726. Illumina WGS data used in this study can be found under the NCBI BioProject accession PRJNA308208. Source code used for analyses is publically available on GitHub (https://github.com/mu-feline-genome/Felis_catus_9.0_analysis).

## Authors’ contributions

Conceptualization: L.A.L., W.J.M., and W.C.W. Data Curation: R.M.B., B.W.D., F.H.G.F, Formal analysis: R.M.B., B.W.D., F.H.G.F., W.A.B, K.K., Funding Acquisition: L.A.L., R.M., W.J.M., and W.C.W. Supervision: L.A.L. Validation: L.A.L. Writing (original draft): R.M.B., L.A.L., and W.C.W. Writing (review and editing): all authors.

## Acknowledgements

We appreciate the donation of funding and samples from cat breeders, especially Terri Harris, who also provided the image in Fig. 4a. We appreciate Thomas R. Juba for his technical assistance of the variant validation and genotyping. We thank Susan Brown at Kansas State University for the generation of the BioNano map.

