## Supplemental information for "A new domestic cat genome assembly based on long sequence reads empowers feline genomic medicine and identifies a novel gene for dwarfism"

**Supplemental Table S1**. Representative annotation measures for assembled carnivore genomes^1^.

| **Species genus** | **Assembled version** | **Assembly level** | **Protein coding genes** | **Total ncRNA** | **mRNAs** | **Repeat masked (%)** |
| --- | --- | --- | --- | --- | --- | --- |
| *Felis catus* | Felis catus 9.0 | Chromosome | 19,738 | 11,679 | 54,713 | 42.72 |
| *Felis catus* | Felis catus 8.0 | Chromosome | 20,176 | 11,868 | 53,014 | 44.09 |
| *Canis lupus familiaris* | CanFam3.1 | Chromosome | 20,039 | 11,743 | 58,761 | 42.96 |
| *Canis lupus dingo* | ASM325472v1 | Scaffold | 20,248 | 12,981 | 62,946 | 42.86 |
| *Acinonyx jubatus* | Aci_jub_2 | Scaffold | 19,529 | 11,000 | 56,248 | 42.66 |
| *Ursus arctos horribilis* | ASM358476v1 | Scaffold | 19,848 | 7,061 | 43,155 | 41.01 |
| *Lynx canadensis* | mLynCan4_v1.p | Chromosome | 19,417 | 6,824 | 44,864 | 43.70 |

^1^All species-specific gene annotation metrics derived from the NCBI database.

**Supplemental Table S2:** Variant calling summary statistics

|  | **All variants** | **SNVs^a^** | **Indels** |
| --- | --- | --- | --- |
| Total | 46,600,527 | 39,043,080 (2.09305) | 13,304,140 |
| Biallelic | 43,213,841 | 36,606,107 (2.44973) | 6,607,734 |
| Multiallelic | 3,386,686 | 2,436,973 | 949,713 |

^a^ Numbers in braces represent ts/tv ratio

**Supplemental Table S3**. Truth sensitivity of SNV call set.

| **Probes** | **Successfully remapped probes** | **Array SNVs found in call set** |
| --- | --- | --- |
| 62,897 | 61,258 (97.39%)^a^ | 59,246 (96.72 %)^b^ |

^a^ Percentage of all probes

^b^ Percentage of successfully remapped probes

**Supplemental Table S4.** SNV classification by minor allele frequency in domestic cats.

| **Consequence** | **Impact** | **MAF < 1%** | **1% < MAF <10%*** | **MAF >10%** | **Total** |
| --- | --- | --- | --- | --- | --- |
| Stop gained | LoF | 212 | 334 | 292 | 838 |
| Start lost | LoF | 40 | 76 | 119 | 235 |
| Stop lost | Lof | 9 | 32 | 63 | 104 |
| Missense variant | Missense | 16476 | 29250 | 31936 | 77662 |
| Synonymous variant | Synonymous | 21812 | 47200 | 59734 | 128746 |
| Stop retained variant | Synonymous | 12 | 38 | 49 | 99 |
| Combined |  | 38562 | 76928 | 92195 | 207685 |

* MAF < 10% refers to an allele count equal to or less than 10 alleles.

**Supplemental** **Table S5**. Feline LoF singletons in human genes under strong constraint.

| **SNV location** | **Consequence** | **SYMBOL** | **pLI** | **Individual ID** | **Supported by NCBI CDS annotation** | **Disease** | **Status** |
| --- | --- | --- | --- | --- | --- | --- | --- |
| chrA1:108024733 | stop_gained | *FBN2* | 0.99999 | felCat.Fcat19725.Fenrisulfr | No | Hypotrichia | affected |
| chrA1:116653102 | stop_gained | *FAM13B* | 0.92003 | felCat.Fcat19194.Pudge | Yes | Ectodermal dysplasia | affected |
| chrA1:192824838 | stop_gained | *CYFIP2* | 1 | felCat.Fcat20406.Gannon | Yes | stones | affected |
| chrA1:195100556 | stop_gained | *LARP1* | 1 | felCat.SPF19984.SPFHarlem | No |  |  |
| chrA1:216234193 | stop_gained | *PDZD2* | 0.99156 | felCat.Fcat18579.Madagascar | No |  |  |
| chrA1:216236592 | stop_gained | *PDZD2* | 0.99156 | felCat.Fcat11849.Iraq | No |  |  |
| chrB1:77305060 | stop_gained | *FBXW7* | 0.99984 | felCat.Fcat5012.Colorado | Yes | Lymphoma | carrier |
| chrB4:41749665 | stop_gained | *ZNF384* | 0.99872 | felCat.S792.Haku | No |  |  |
| chrB4:86010265 | stop_gained | *KIF5A* | 0.9999 | felCat.SPF19984.SPFHarlem | Yes |  |  |
| chrC1:14006692 | stop_gained | *UBR4* | 1 | felCat.Fcat17994.Camila | No | Hydrocephalus | carrier |
| chrC1:14006828 | start_lost | *UBR4* | 1 | felCat.Fcat18849.Iowa | No |  |  |
| chrC2:69450306 | stop_gained | *MYLK* | 0.96358 | felCat.Fcat20425.Rocket | No | Hypokalemia | affected |
| chrD1:108097571 | stop_gained | *INCENP* | 0.98884 | felCat.Fcat18801.Italy | No |  |  |
| chrD2:63980875 | stop_gained | *SH3PXD2A* | 0.99821 | felCat.CR1397.Isabella | Yes | Infectious peritonitis | affected |
| chrD4:16547403 | stop_gained | *RORB* | 0.99959 | felCat.Fcat18528.Denmark | Yes |  |  |
| chrD4:75416950 | stop_gained | *PRPF4* | 0.99926 | felCat.Fcat18528.Denmark | Yes |  |  |

**Supplemental Table S6.** GATK variant filtering criteria.

| **Filter name** | **SNV GATK** | **SNV Strict** | **Indel GATK** | **Indel Strict** |
| --- | --- | --- | --- | --- |
| QD | X < 2.0 | X < 8.0 | X < 2.0 | X < 8.0 |
| FS | X > 60.0 | X > 20.0 | X > 200.0 | X > 20.0 |
| SOR | X > 3.0 | X > 2.0 | X > 10.0 | X > 2.0 |
| ReadPosRankSum | X < -8.0 | X < -2.0 \|\| X > 2.0 | X < -20.0 | X < -2.0 \|\| X > 2.0 |
| MQ | X < 40.0 | X < 55.0 | NA | X < 55.0 |
| MQRankSum | X < -12.5 | X < -0.2 \|\| X > 0.2 | NA | X < -0.2 \|\| X > 0.2 |

Filters are as follows: variant quality by read depth (QD), fisher test for strand (FS), strand odds ratio (SOR), read position rank sum test (ReadPosRankSum), mapping quality (MQ), and mapping quality rank sum (MQRankSum).

**Supplemental** **Table S7**. PCR Primers for the genotyping of feline disproportionate dwarfism.

| **Number on Fig S1** | **Primer** | **Sequence 5’ 🡪 3’** |
| --- | --- | --- |
| 1 | UGDH_mid_R | TGGAGATGTGCACCTTCATC |
| 2 | UGDH_mid_F | GGCGTAAACACATTTTCTTGC |
| 3 | UGDH_del_R | CGGCATACAAGTCAGCCTTC |
| 4 | UGDH_down_R | GGGCAAAATTGGGGACTAAC |
| 5 | UGDH_up_F | CAGTGTGTGGCATAGGCTTC |

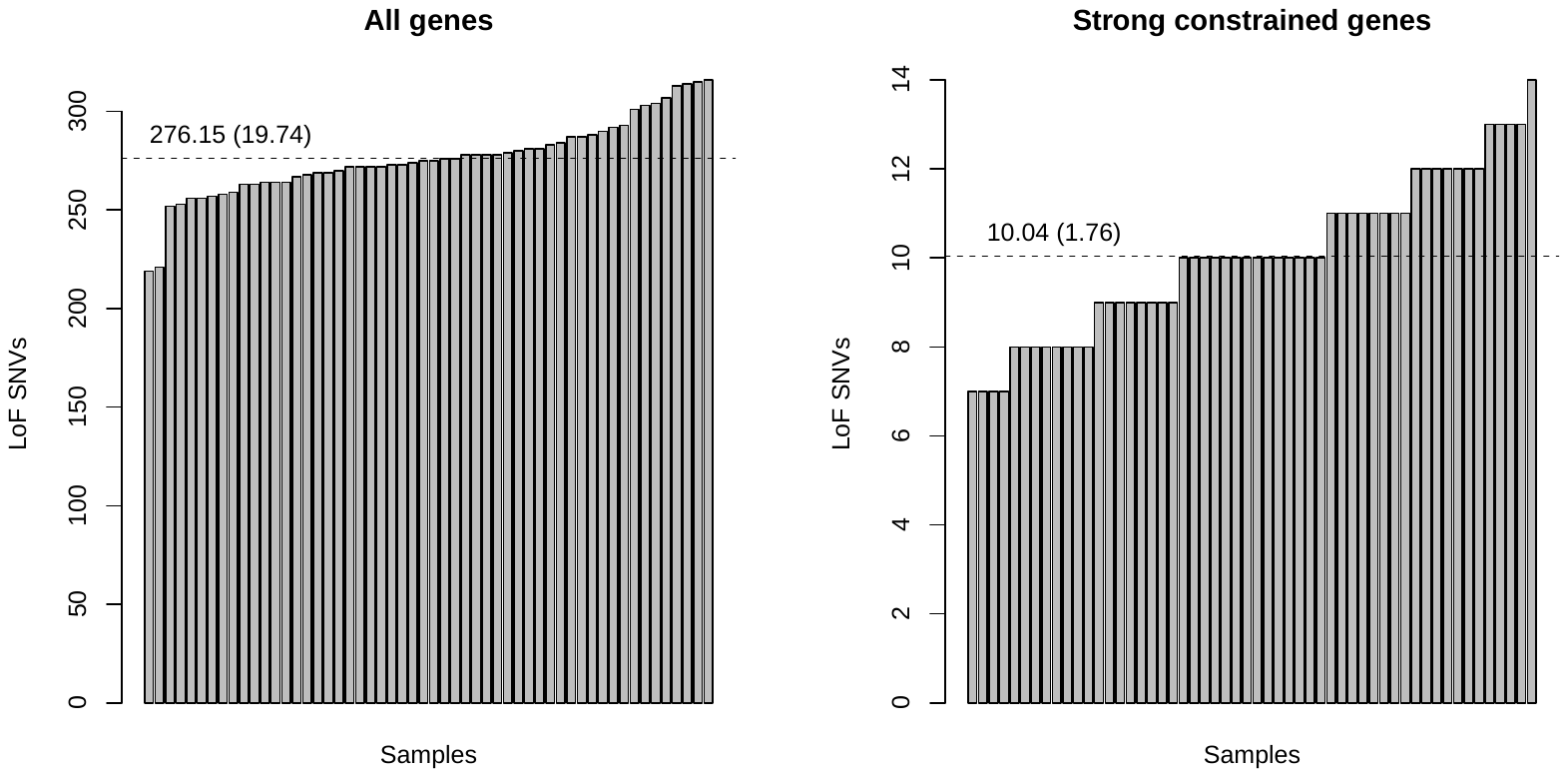

**Supplemental Figure S1**. LoF SNVs per individual in all genes and strong constrained genes. Dotted line shows the mean value, which is also stated above along with standard deviation in braces.

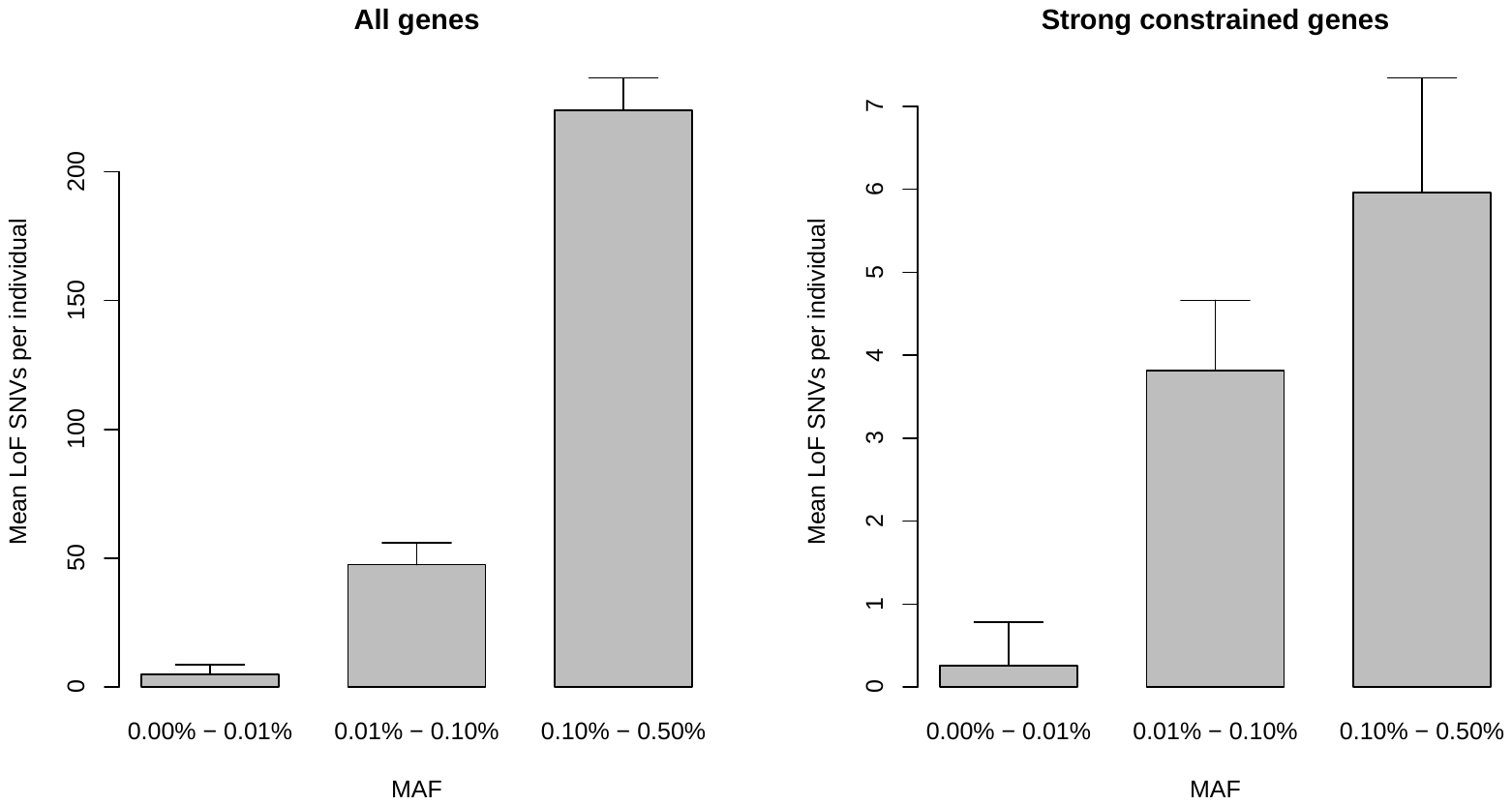

**Supplemental Figure S2**. Mean LoF SNVs per individual grouped by minor allele frequency in all genes and strong constrained genes. Error bars represent 1 standard deviation.

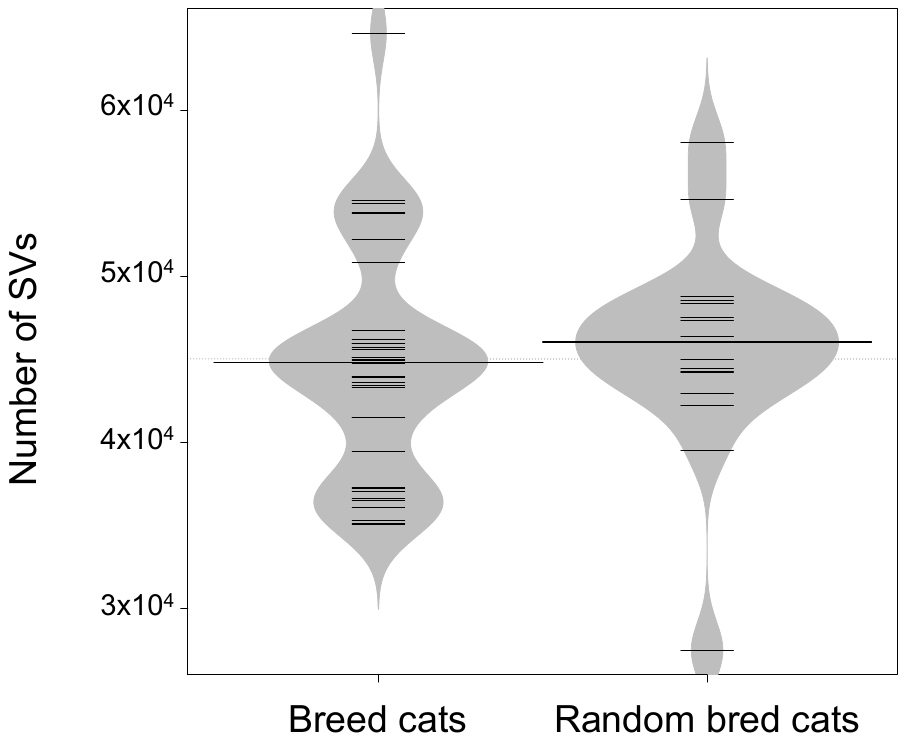

**Supplemental Figure S3**. Number of SVs found in breed cats and random bred cats.

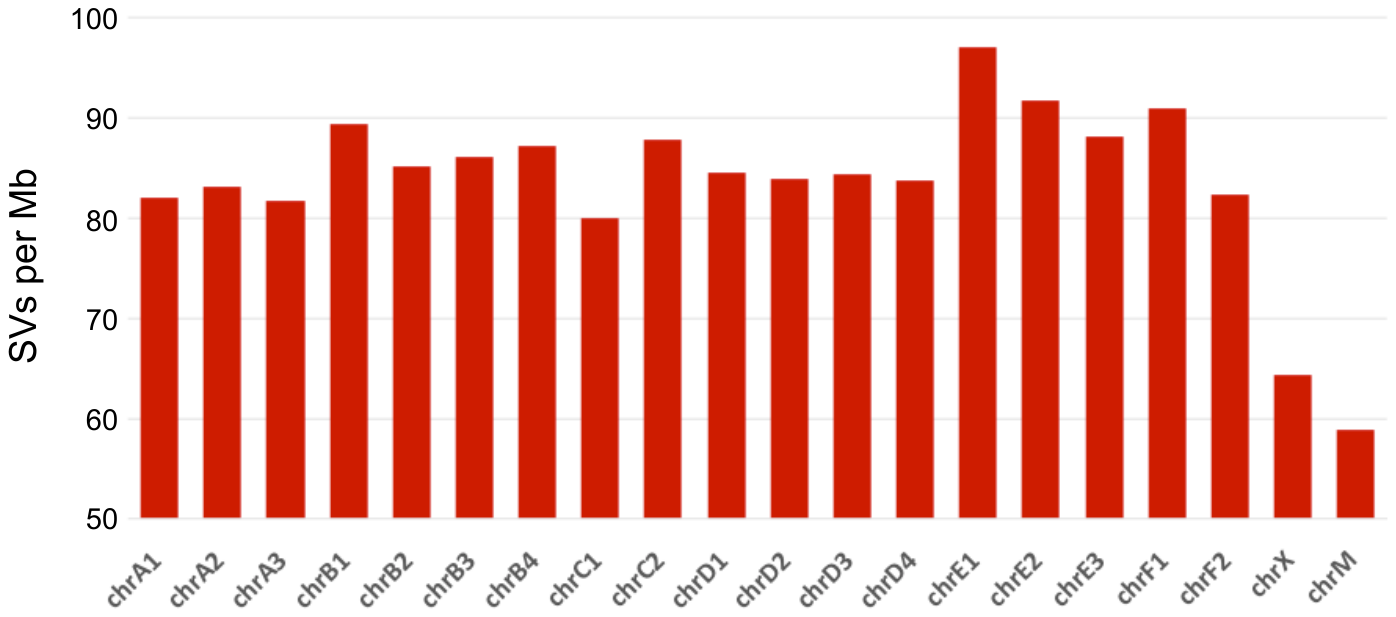

**Supplemental Figure S4**. SV frequency per chromosome.

**Supplemental Figure S5.** Discordant reads overlapping *UGDH* are unique to unrelated affected cats. Unrelated affected cats are felCat.19050.Mouse, felCat.19060.Gwenivere, and felCat.19067.Princess. felCat.17799.Cali is an unaffected normal control cat. Coverage across the control cat is relatively uniform, while the affected cats show decreased coverage over the final exon of UGDH marking a heterozygous deletion. Discordant reads that span beyond the deletion show sequence into the deleted region shares homology with *UGDH* exon 8.

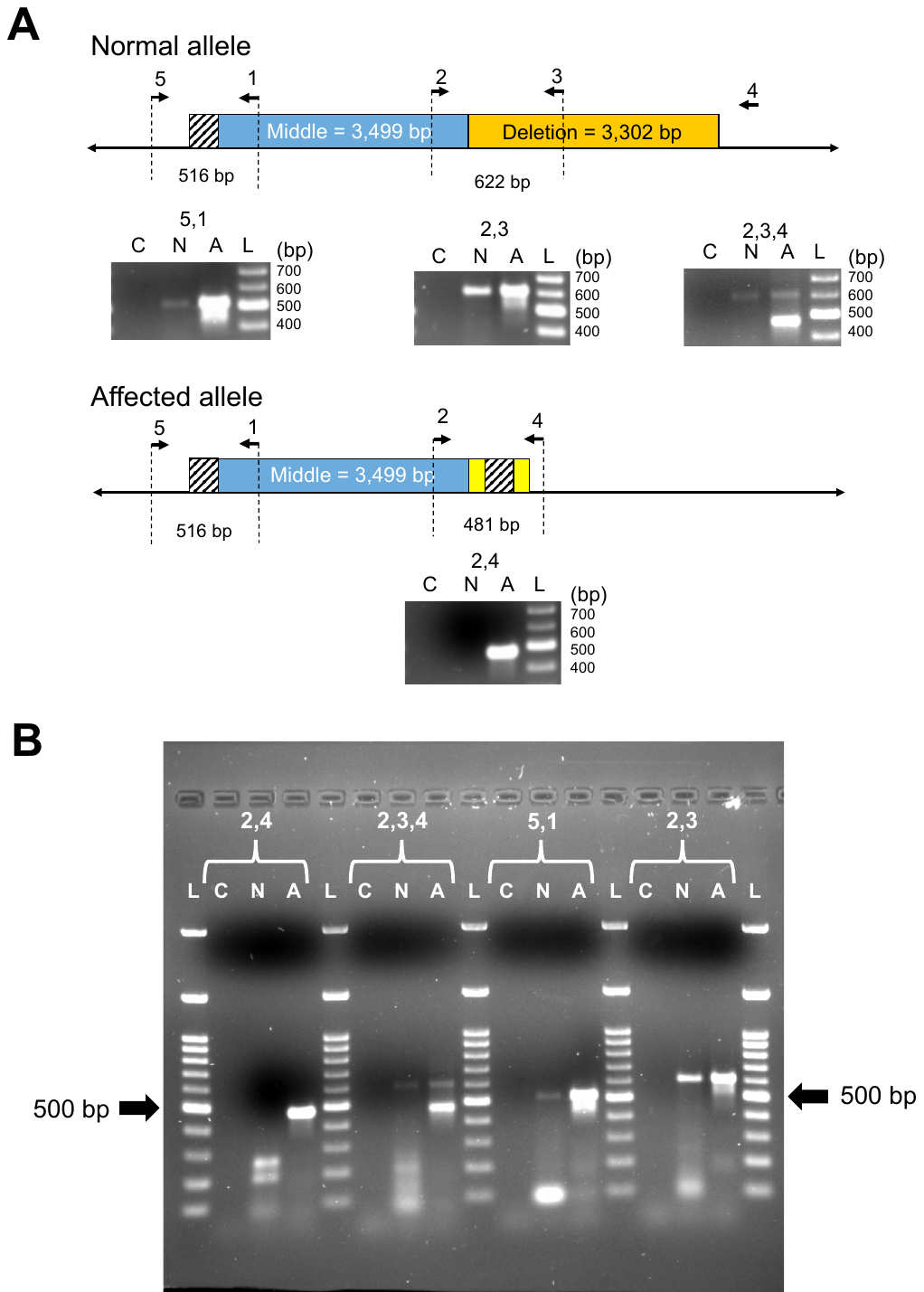

**Supplemental Figure S6**. Breakpoints validated for dwarfism SV in *UGDH.* **(A)** Arrows represent individual primers and predicted band sizes. Gel photos show no template control (C), normal sample (N), affected sample (A), and 100 bp PLUS™ DNA Ladder (Gold Biotechnology, Inc., St. Louis, MO) (L). Ladder sizes are shown to the right of each gel image in bp. Above each gel image is the primers that were used to generate the band in each sample. Band sizes were consistent with predicted breakpoint lengths. Hashed square is 49 bp segment that shares homology with exon 8, it is consistent with a duplication and insertion into deleted region. Yellow boxes represent sequence of unknown origin found in affected allele. The deletion is absent from the affected allele, allowing primer four to produce an amplicon with primer 2. All dwarf samples analyzed were heterozygous for the affected allele. Primers 1 – UDGH_mid_R, 2 – UDGH_mid_F, 3 – UDGH_del_R, 4 – UDGH_down_R, 5 – UDGH_up_F. **(B)** Full gel image for primer combinations described in A.

**Supplemental Figure S7.** Histology of control and dwarf cartilage plates. H&E and toluidine blue histologic samples of the distal radius epiphyseal cartilage plate from a normal neonatal kitten and age-matched dwarf kittens. For normal control kitten, H&E shows chondrocytes in the growth plate exhibit a regular columnar arrangement and are organized into a zone of reserve cells, a zone of proliferation and a zone of hypertrophy and a zone of provisional calcification. For the same kitten toluidine blue stain shows physeal cartilage contains abundant proteoglycan as shown by its metachromasia. In dwarf samples H&E staining consistently shows chondrocytes in the growth plate exhibit an irregular columnar arrangement. For four of the six dwarf samples, toluidine blue staining shows by its metachromasia that dwarf cat physeal cartilage contains lessor amounts of proteoglycans.
