## Supplementary material for "A new domestic cat genome assembly based on long sequence reads empowers feline genomic medicine and identifies a novel gene for dwarfism": S1 Data

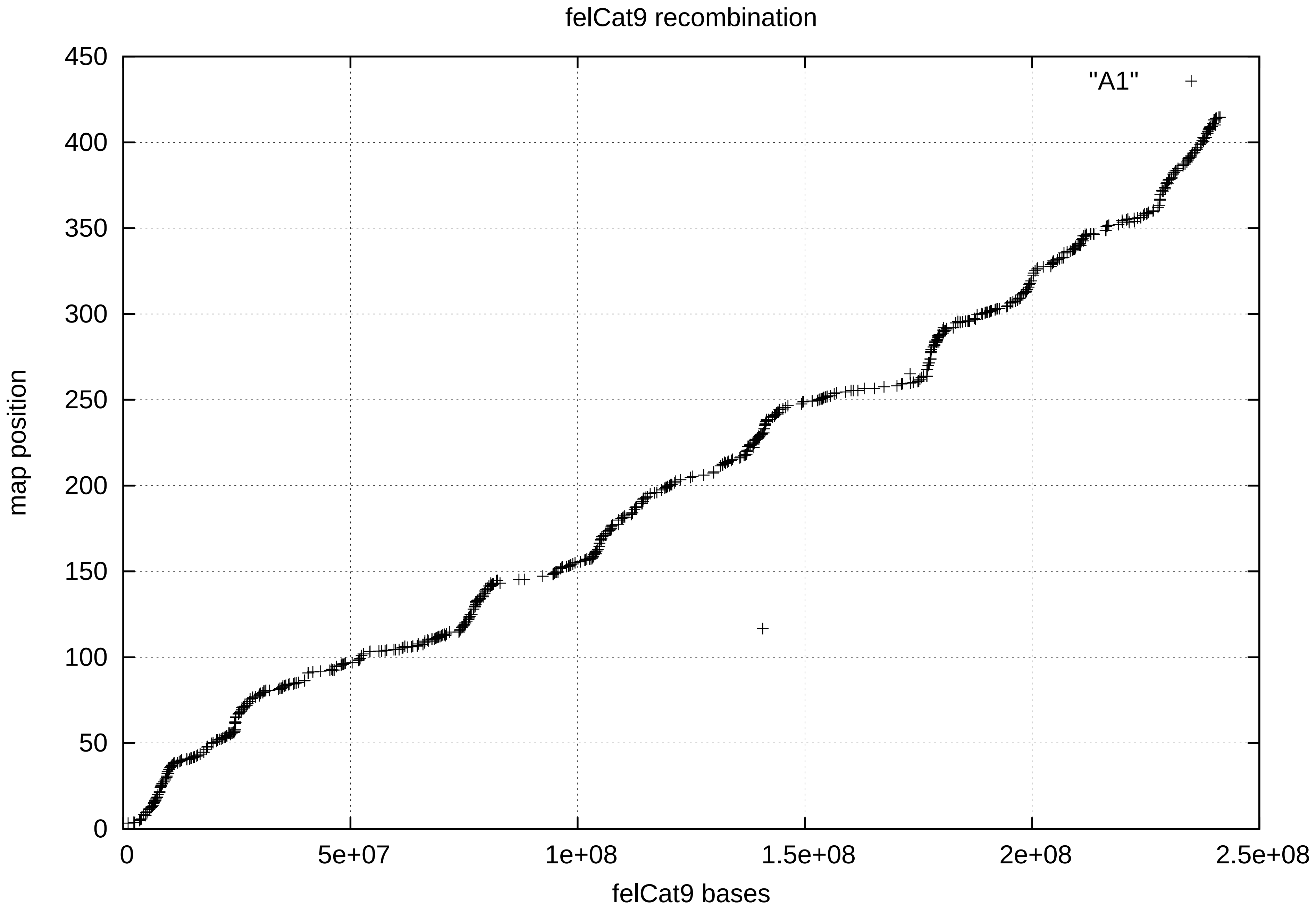

felCat9 recombination
"A1"
map position
felCat9 bases
0
50
100
150
200
250
300
350
400
450
0
5e+07
1e+08
1.5e+08
2e+08
2.5e+08

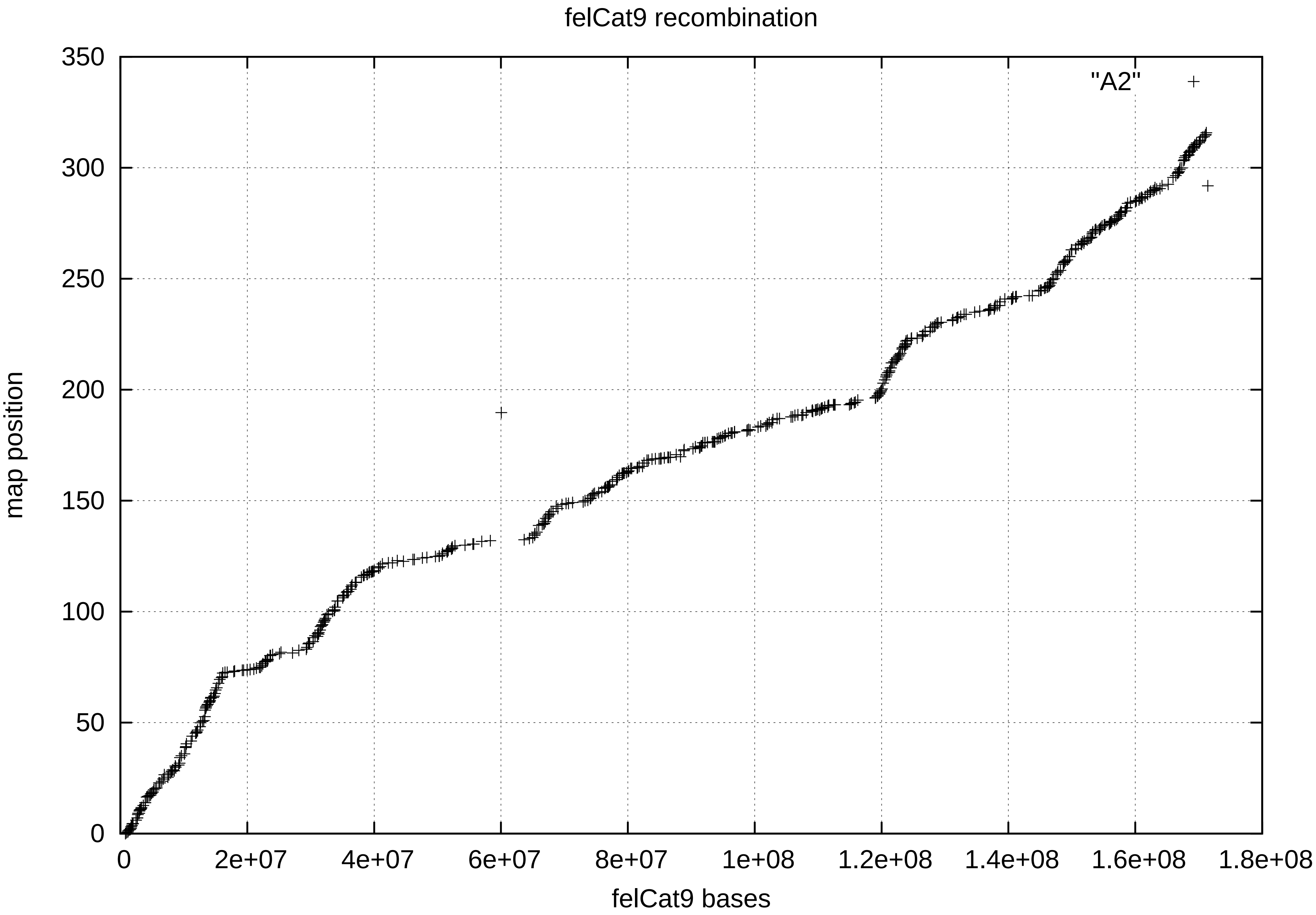
felCat9 recombination
"A2"
map position
felCat9 bases
0
50
100
150
200
250
300
350
0
2e+07
4e+07
6e+07
8e+07
1e+08
1.2e+08
1.4e+08
1.6e+08
1.8e+08

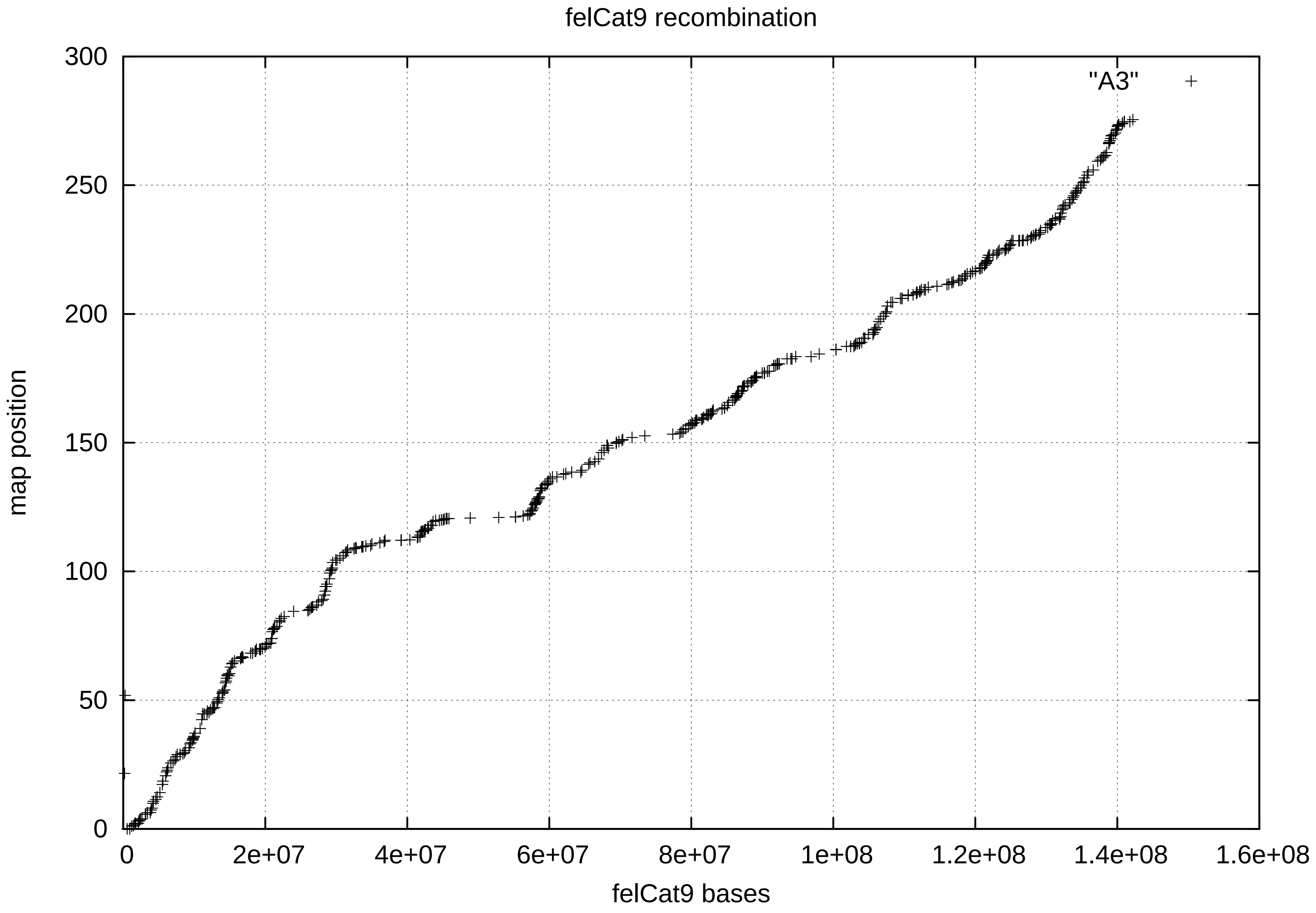

felCat9 recombination
"A3"
map position
felCat9 bases
0
50
100
150
200
250
300
0
2e+07
4e+07
6e+07
8e+07
1e+08
1.2e+08
1.4e+08
1.6e+08

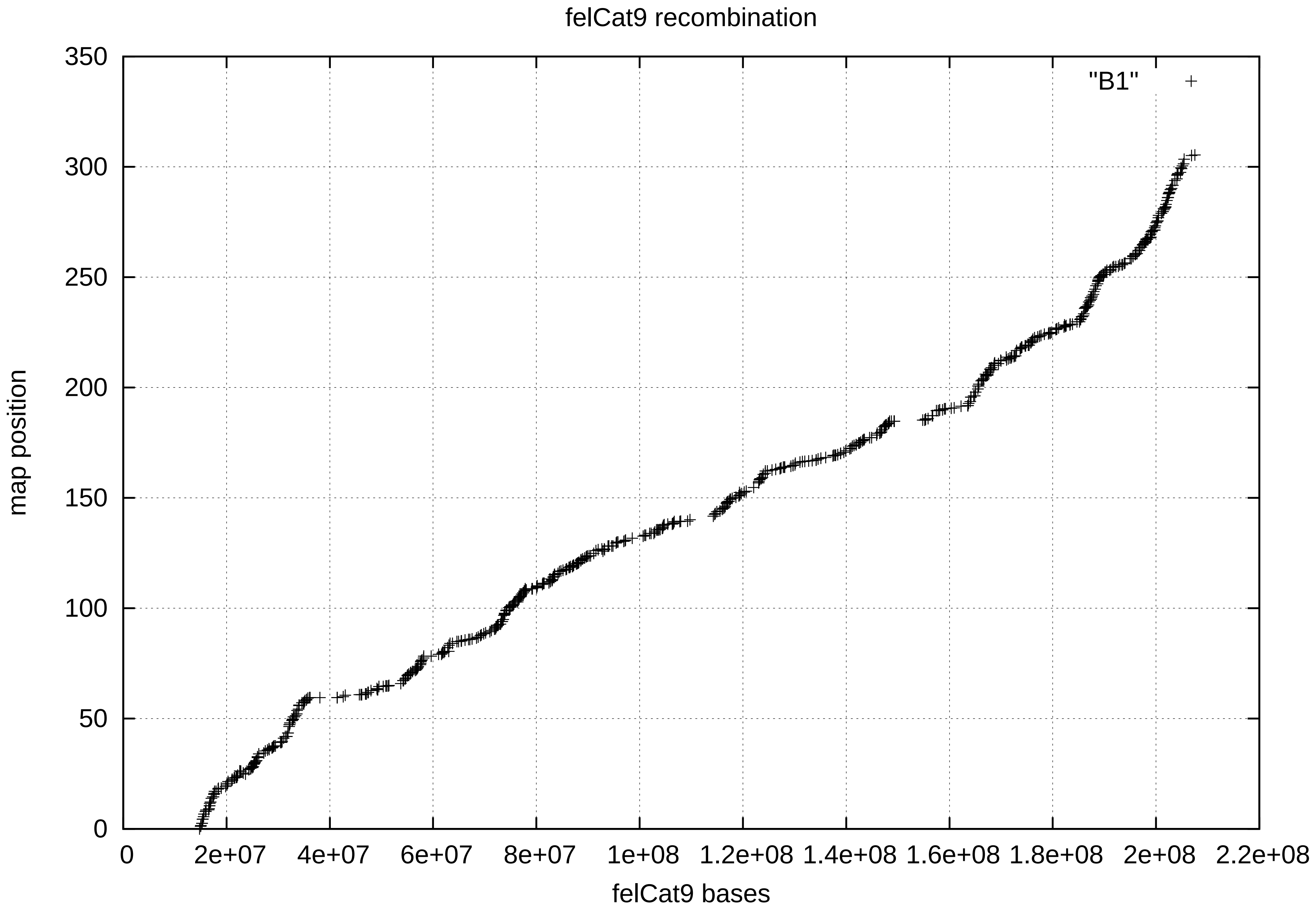

felCat9 recombination
"B1"
map position
felCat9 bases
0
50
100
150
200
250
300
350
0
2e+07
4e+07
6e+07
8e+07
1e+08
1.2e+08
1.4e+08
1.6e+08
1.8e+08
2e+08
2.2e+08

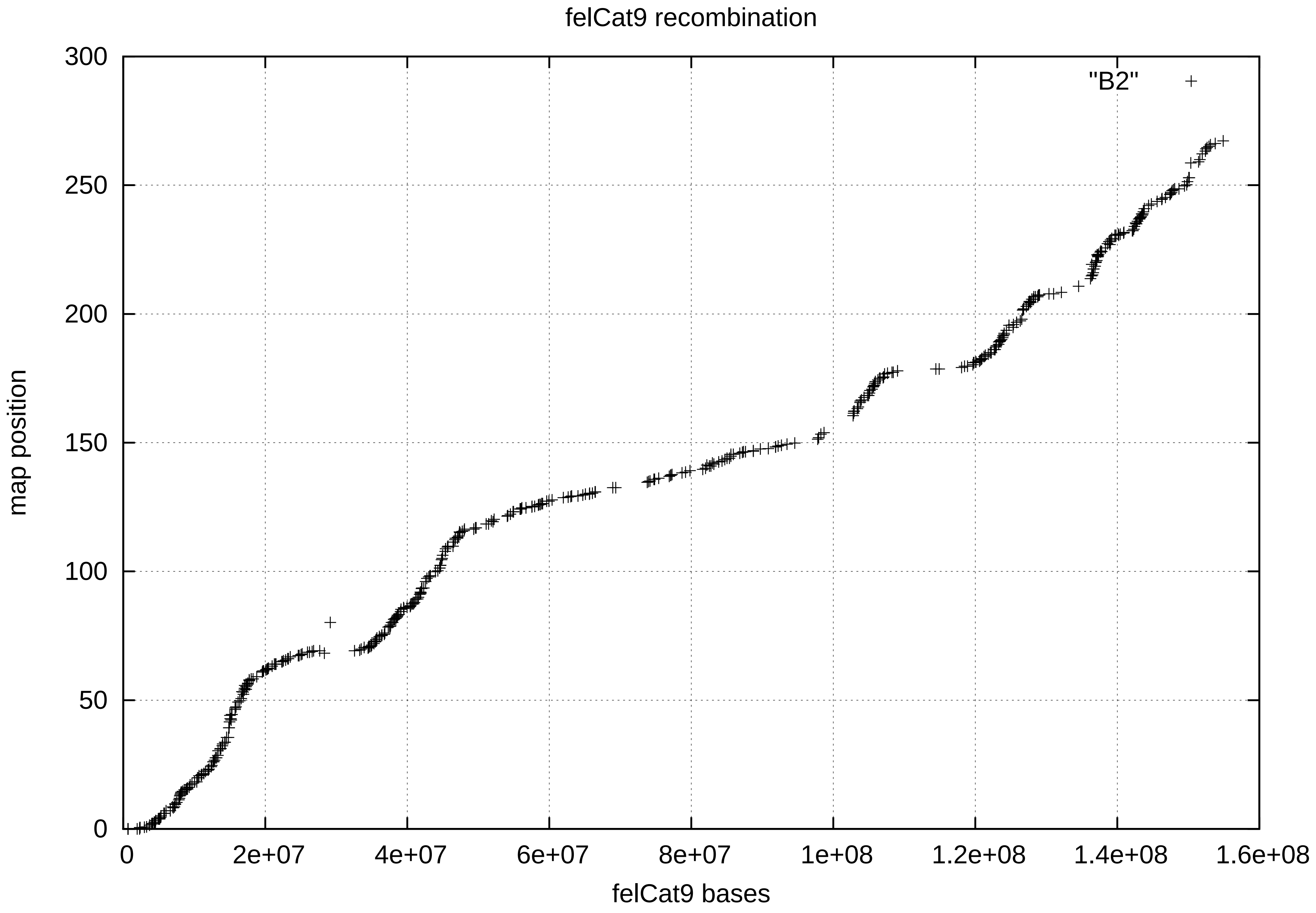

felCat9 recombination
"B2"
map position
felCat9 bases
300
250
200
150
100
50
0
0
2e+07
4e+07
6e+07
8e+07
1e+08
1.2e+08
1.4e+08
1.6e+08

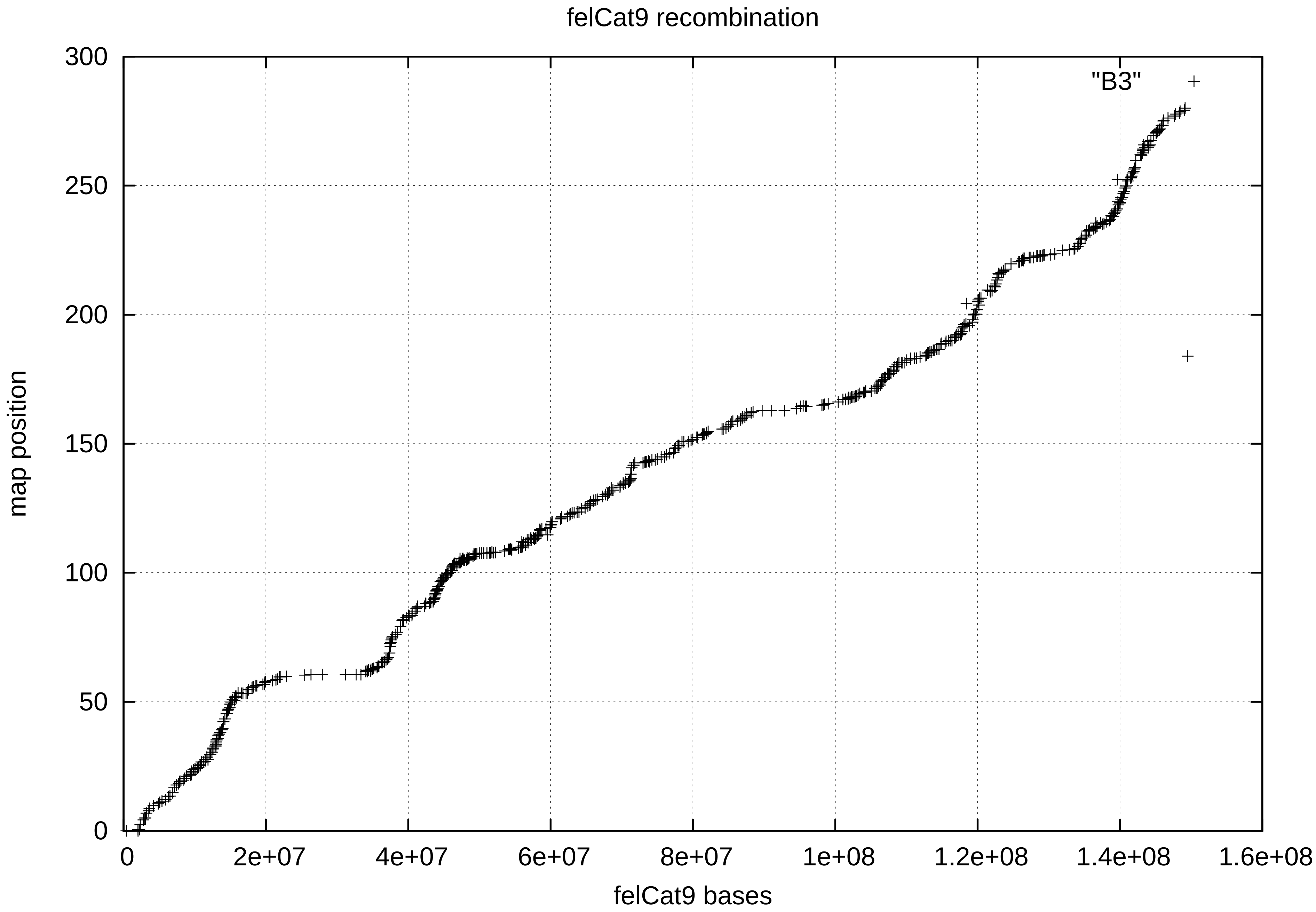
felCat9 recombination
"B3"
map position
felCat9 bases
0
50
100
150
200
250
300
0
2e+07
4e+07
6e+07
8e+07
1e+08
1.2e+08
1.4e+08
1.6e+08

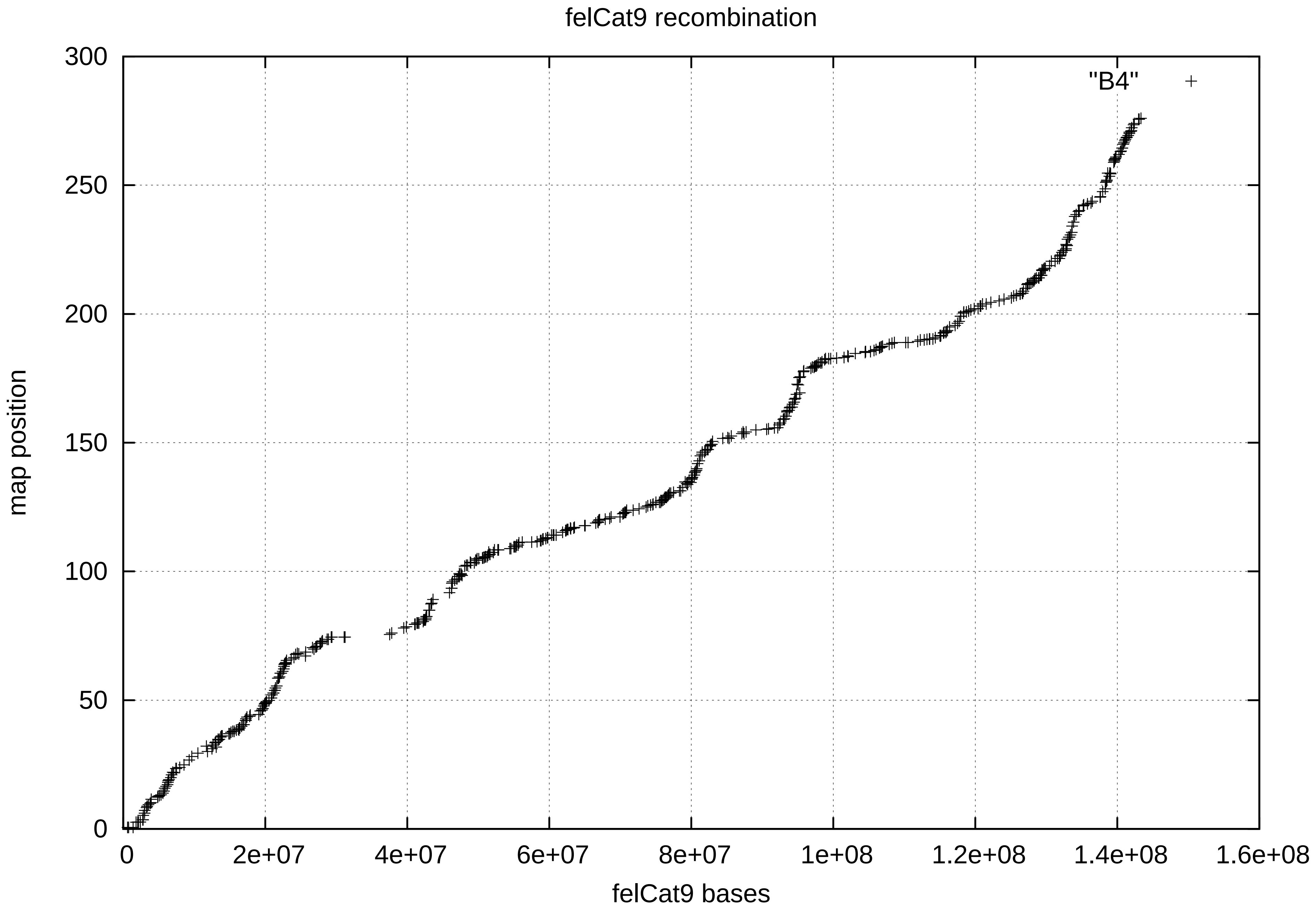

felCat9 recombination
"B4"
map position
felCat9 bases
0
50
100
150
200
250
300
0
2e+07
4e+07
6e+07
8e+07
1e+08
1.2e+08
1.4e+08
1.6e+08

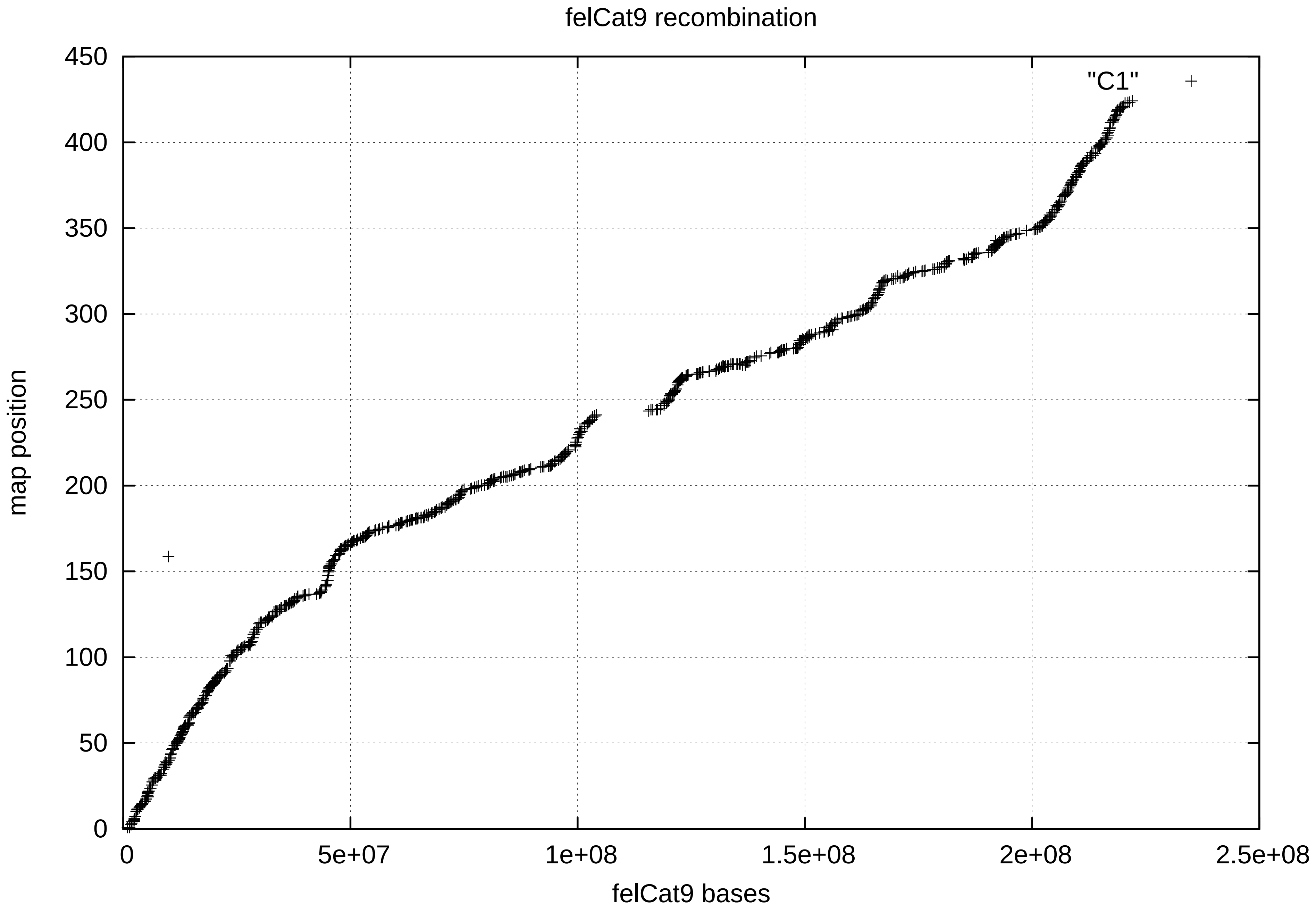

felCat9 recombination
"C1"
map position
felCat9 bases
0
50
100
150
200
250
300
350
400
450
0
5e+07
1e+08
1.5e+08
2e+08
2.5e+08

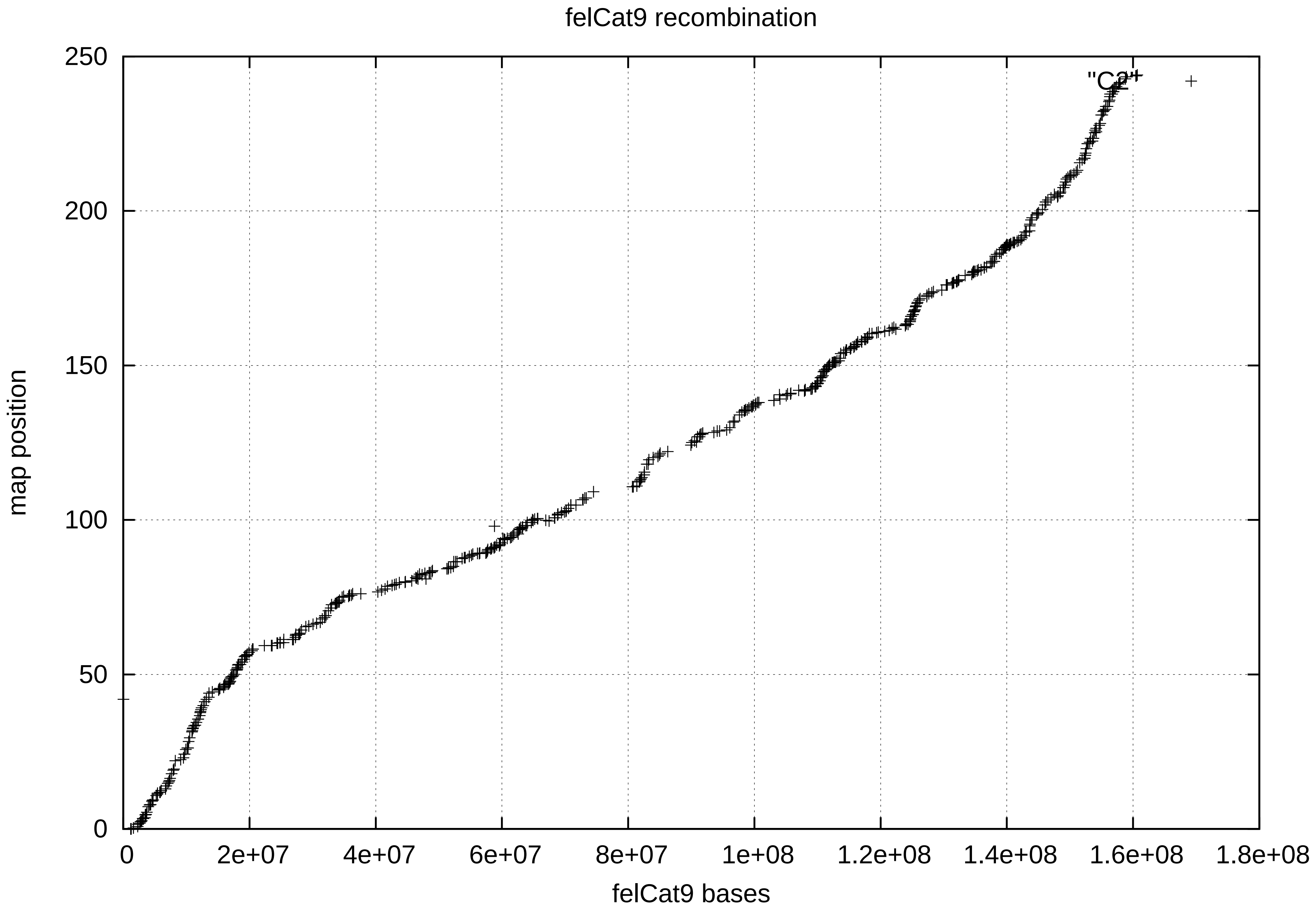

felCat9 recombination
"C2"
map position
0
50
100
150
200
250
0
2e+07
4e+07
6e+07
8e+07
1e+08
1.2e+08
1.4e+08
1.6e+08
1.8e+08
felCat9 bases

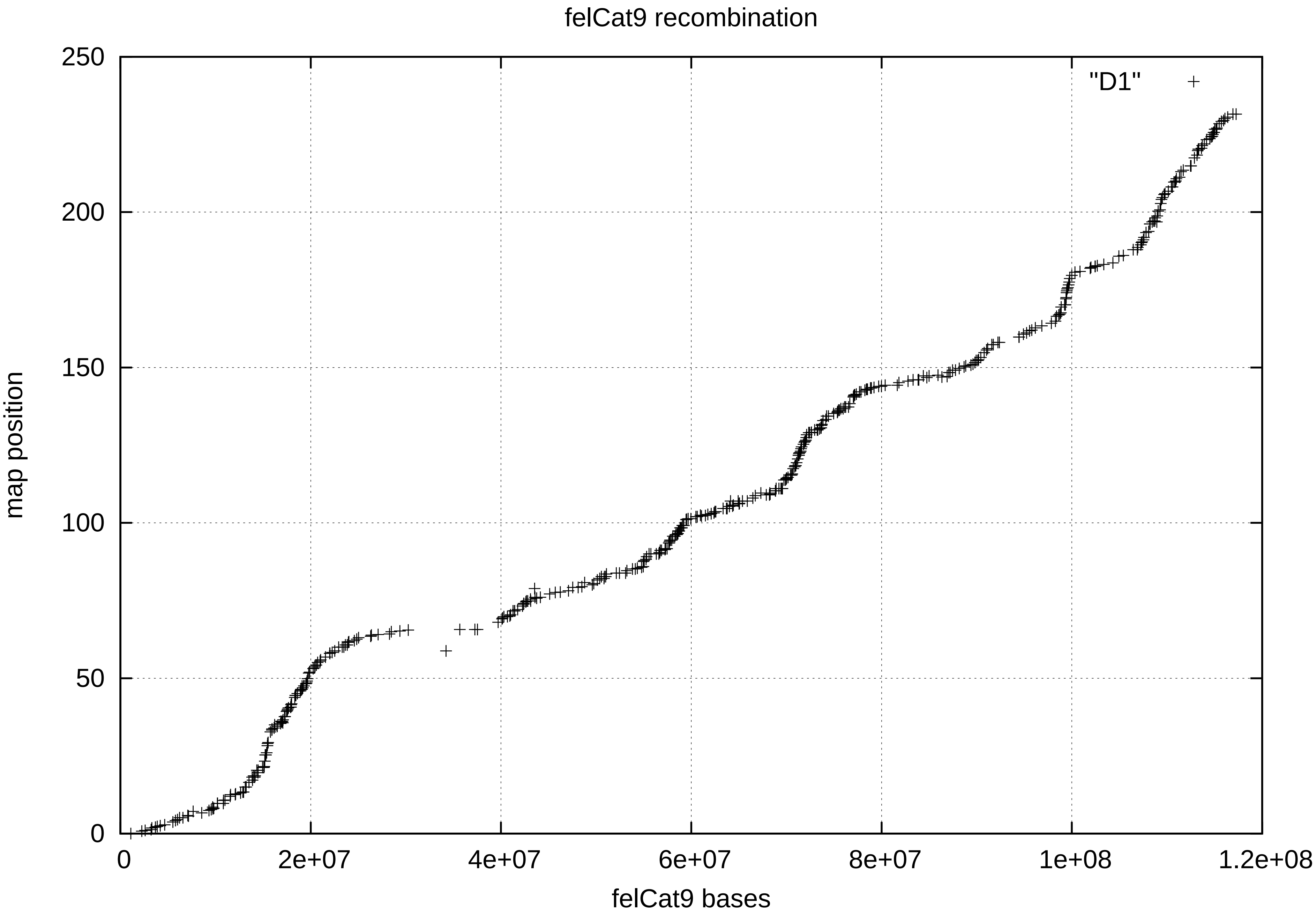

felCat9 recombination
"D1"
map position
felCat9 bases
0
50
100
150
200
250
0
2e+07
4e+07
6e+07
8e+07
1e+08
1.2e+08

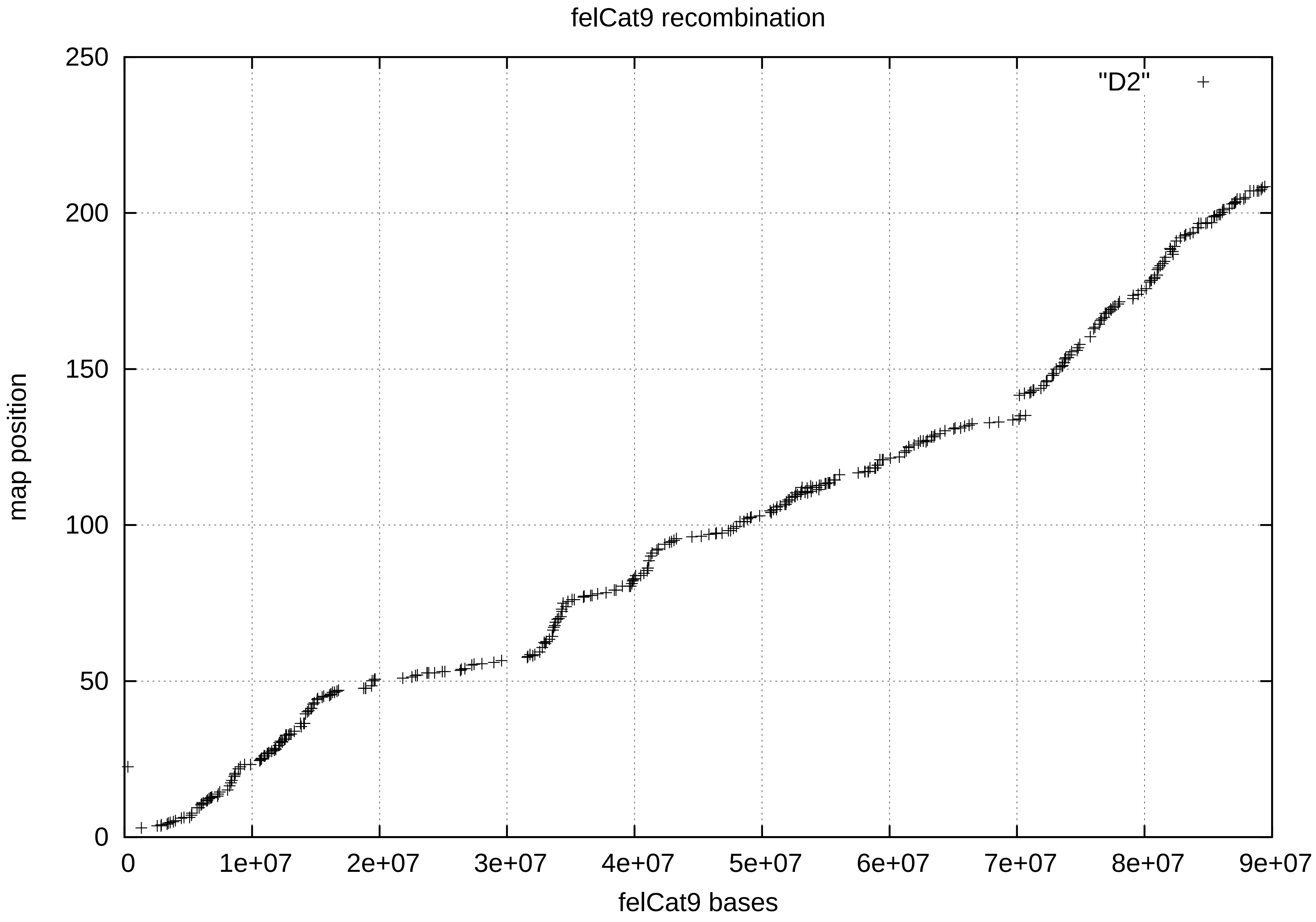

felCat9 recombination
"D2"
250
200
150
100
50
0
map position
0
1e+07
2e+07
3e+07
4e+07
5e+07
6e+07
7e+07
8e+07
9e+07
felCat9 bases

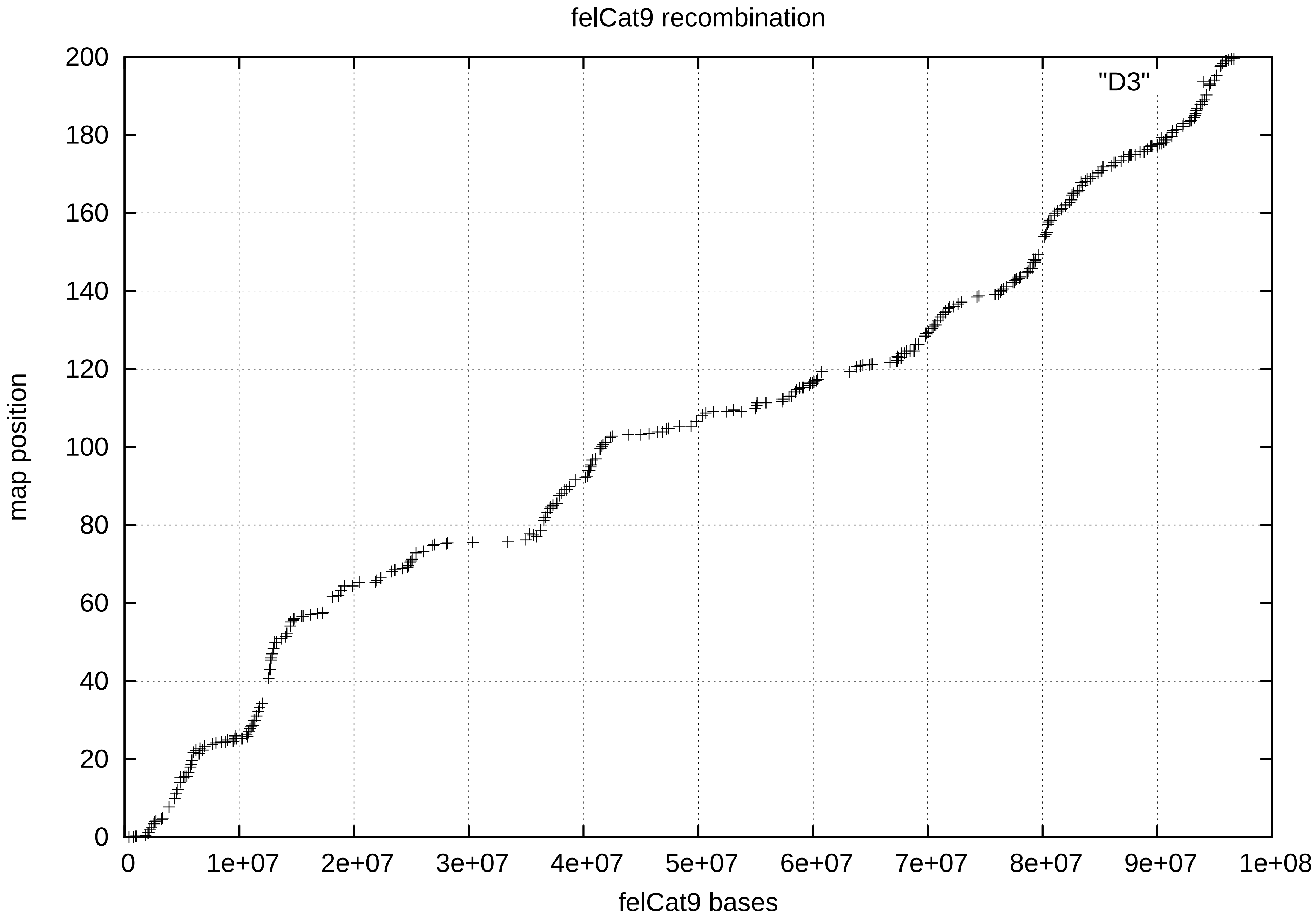
felCat9 recombination
"D3"
map position
0
20
40
60
80
100
120
140
160
180
200
0
1e+07
2e+07
3e+07
4e+07
5e+07
6e+07
7e+07
8e+07
9e+07
1e+08
felCat9 bases

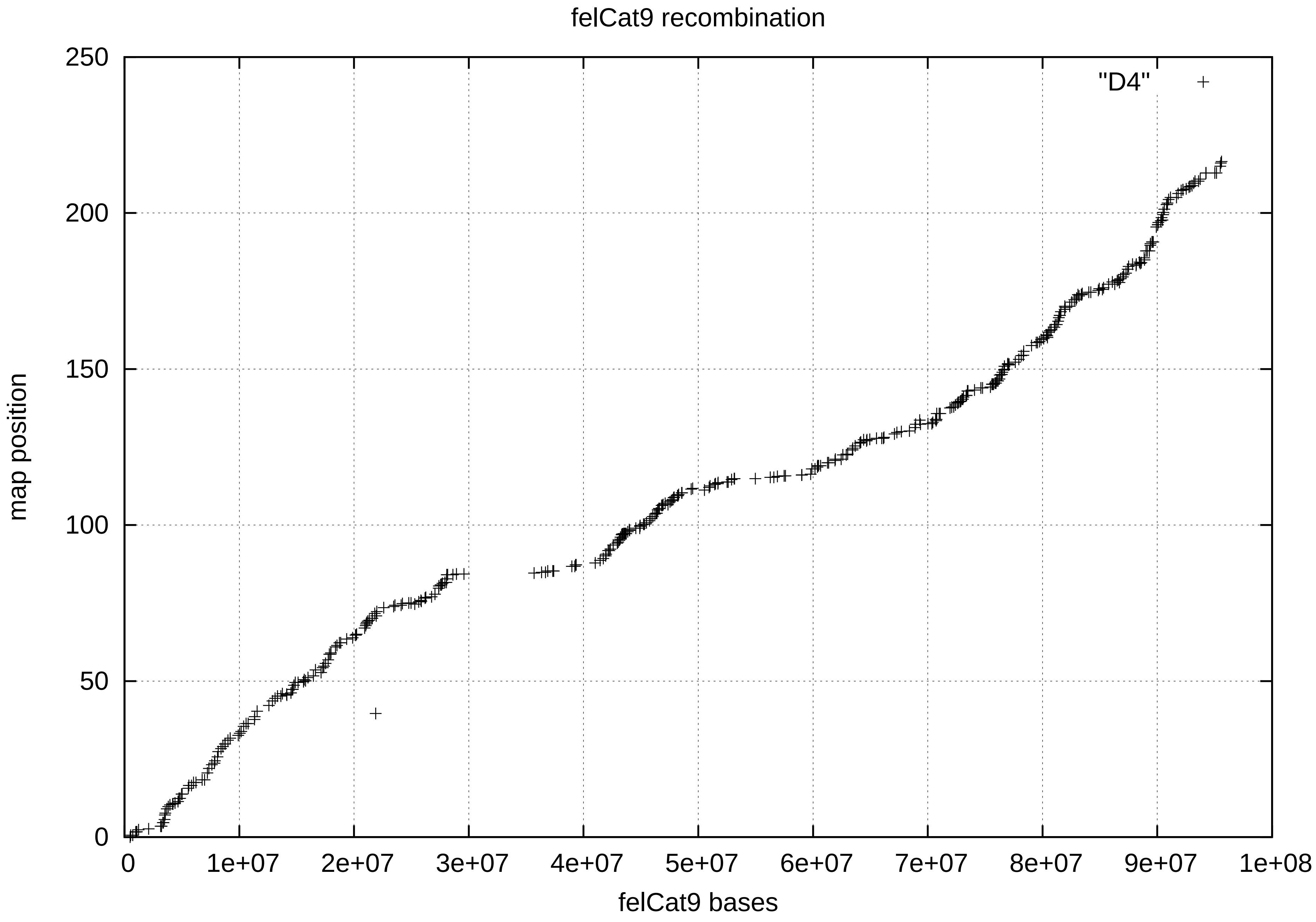
felCat9 recombination
"D4"
250
200
150
100
50
0
map position
0
1e+07
2e+07
3e+07
4e+07
5e+07
6e+07
7e+07
8e+07
9e+07
1e+08
felCat9 bases

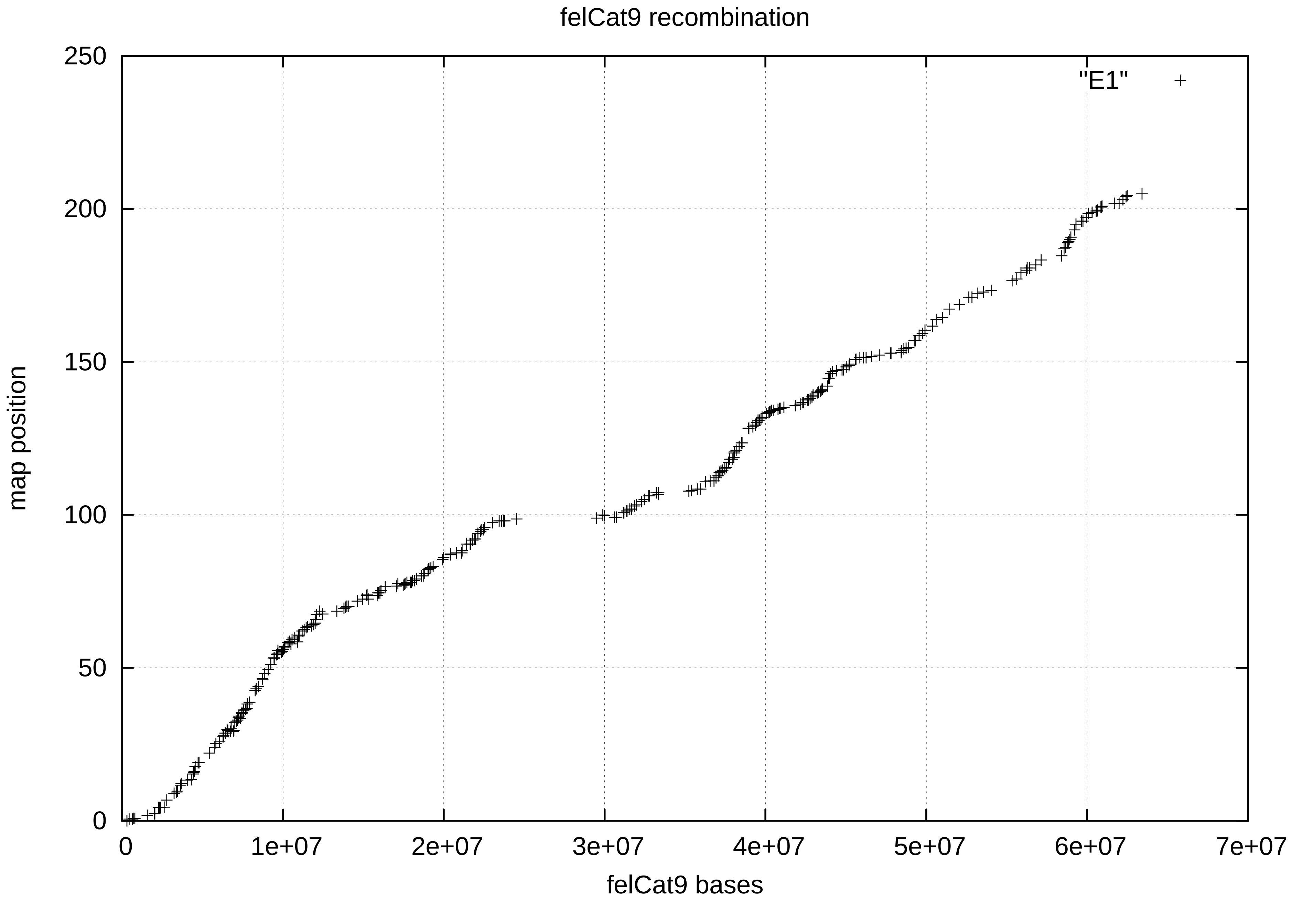
felCat9 recombination
map position
felCat9 bases
"E1"
0
50
100
150
200
250
0
1e+07
2e+07
3e+07
4e+07
5e+07
6e+07
7e+07

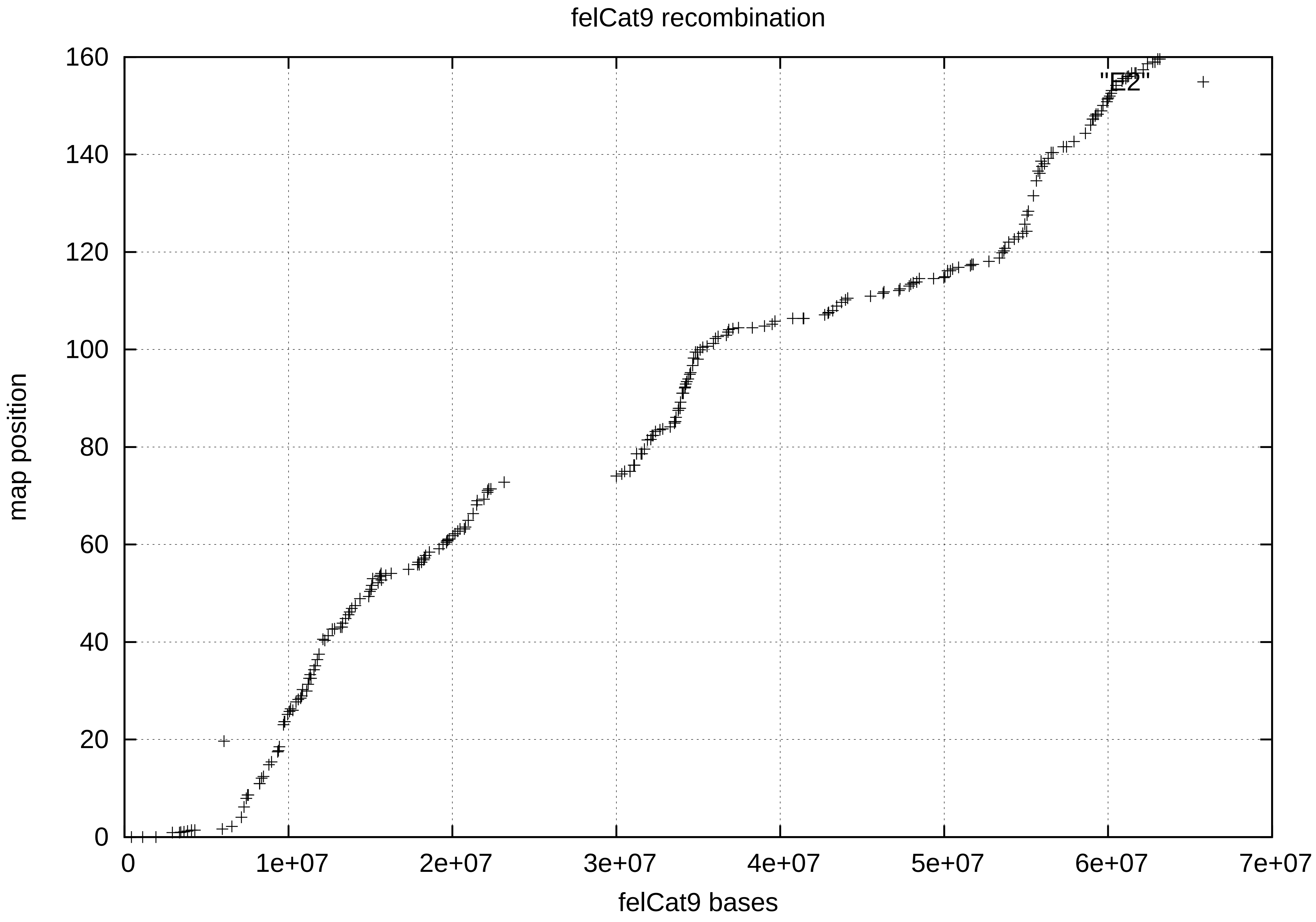

felCat9 recombination
"E2"
160
140
120
100
80
60
40
20
0
map position
0
1e+07
2e+07
3e+07
4e+07
5e+07
6e+07
7e+07
felCat9 bases

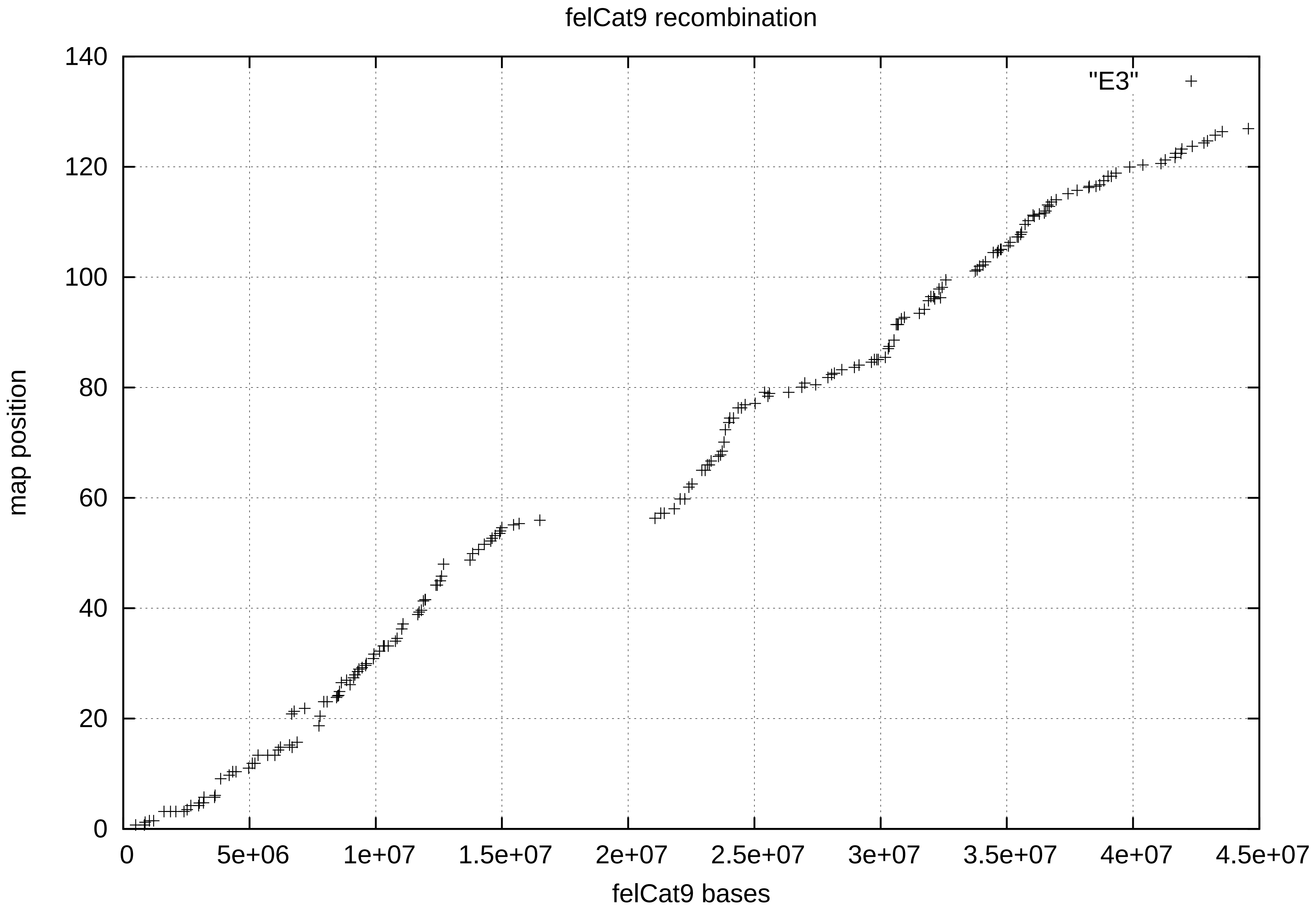

felCat9 recombination
"E3"
map position
felCat9 bases
0
20
40
60
80
100
120
140
0
5e+06
1e+07
1.5e+07
2e+07
2.5e+07
3e+07
3.5e+07
4e+07
4.5e+07

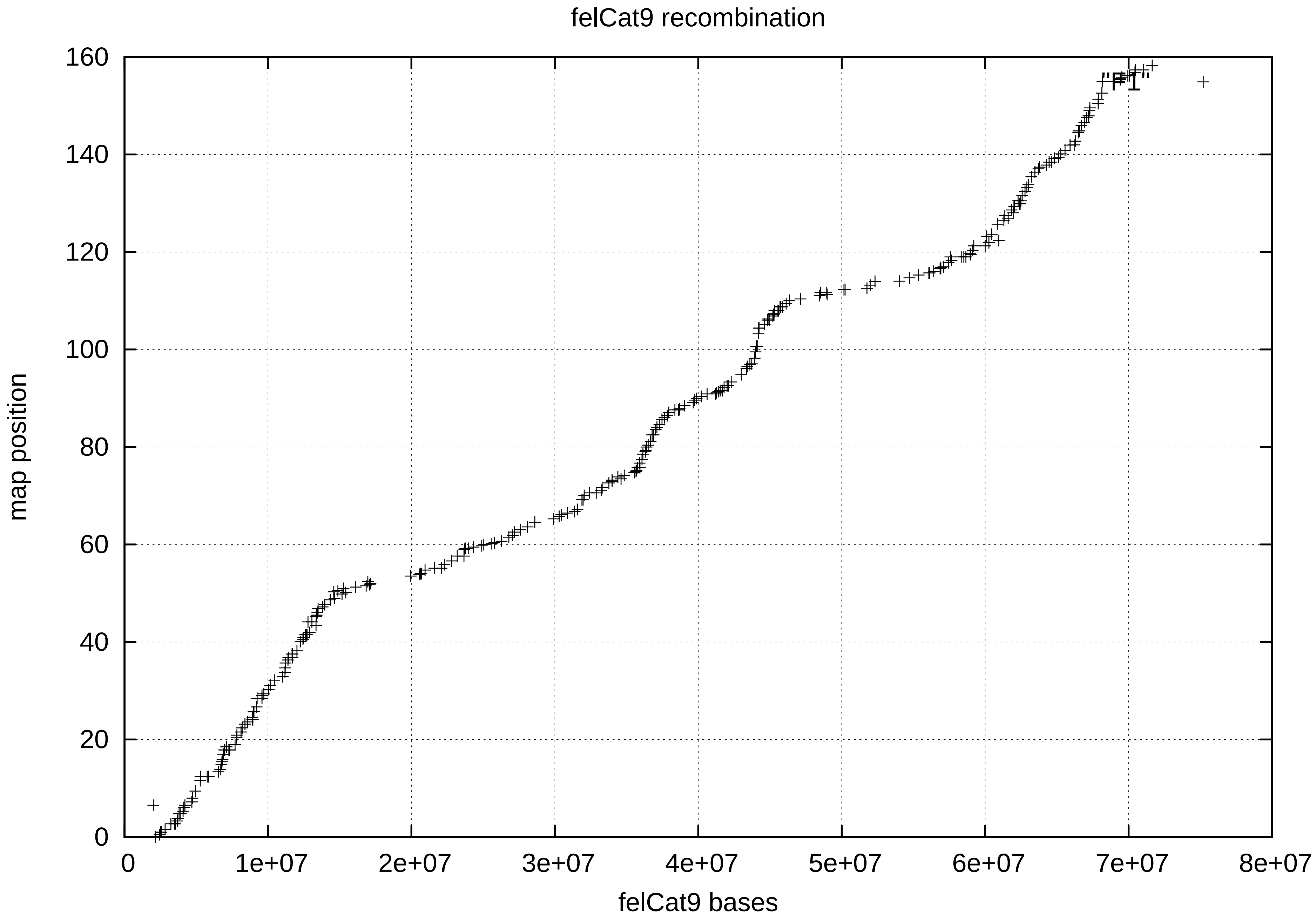
felCat9 recombination
160
140
120
100
80
60
40
20
0
map position
"F1"
0
1e+07
2e+07
3e+07
4e+07
5e+07
6e+07
7e+07
8e+07
felCat9 bases

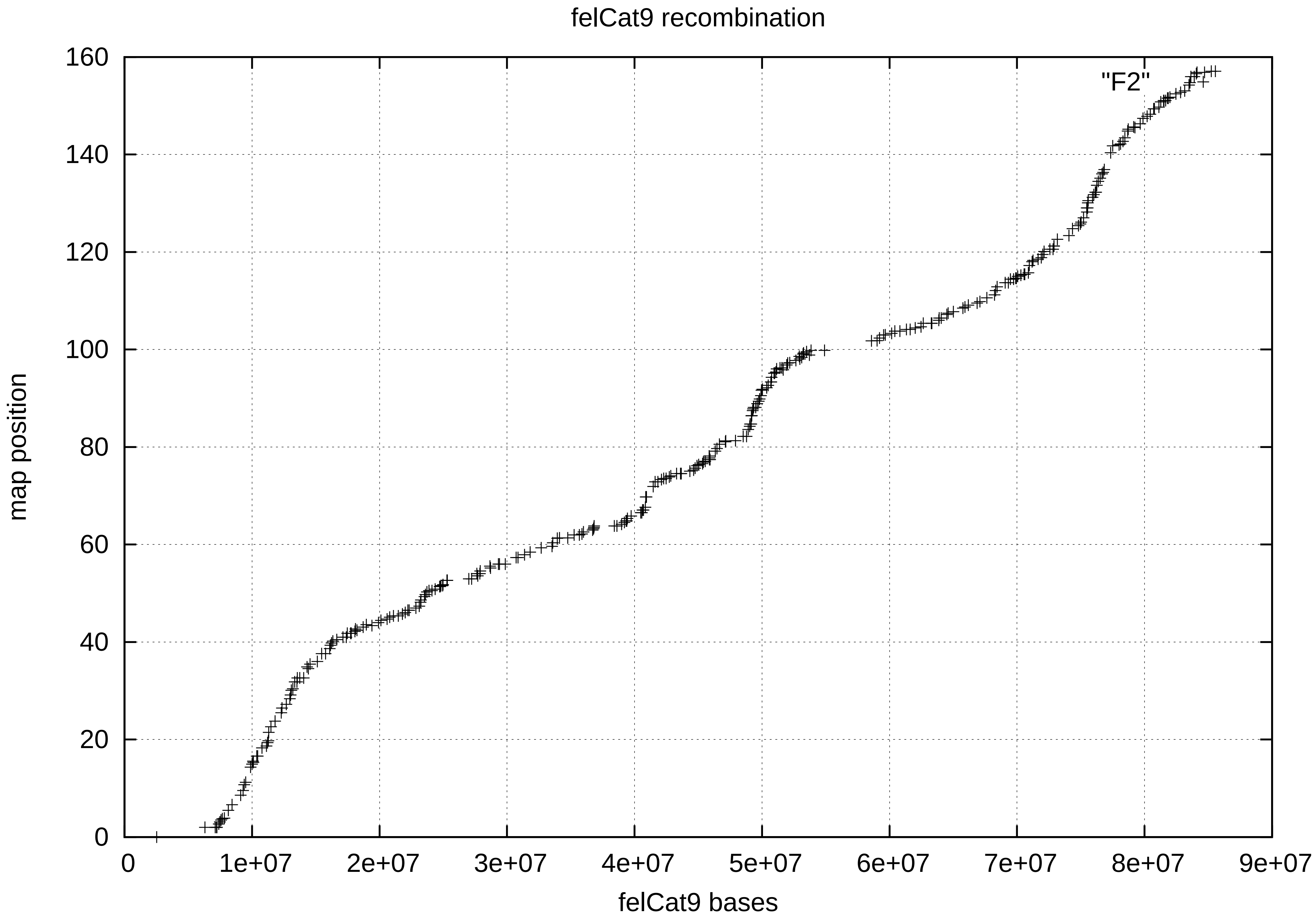
felCat9 recombination
"F2"
map position
0
20
40
60
80
100
120
140
160
0
1e+07
2e+07
3e+07
4e+07
5e+07
6e+07
7e+07
8e+07
9e+07
felCat9 bases

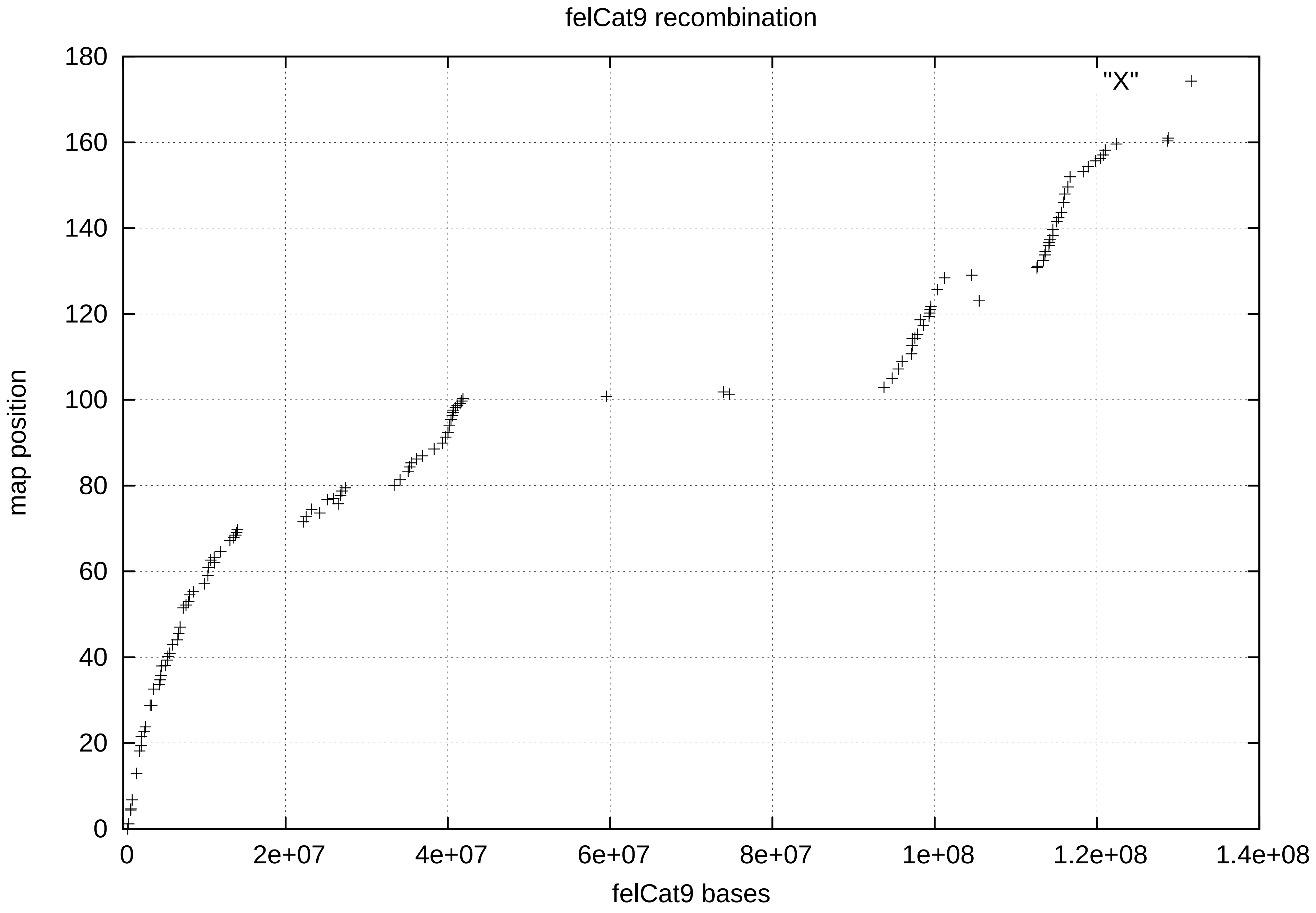
felCat9 recombination
map position
felCat9 bases
"X"
0
20
40
60
80
100
120
140
160
180
0
2e+07
4e+07
6e+07
8e+07
1e+08
1.2e+08
1.4e+08
